# CNest: A Novel Copy Number Association Discovery Method Uncovers 862 New Associations from 200,629 Whole Exome Sequence Datasets in the UK Biobank

**DOI:** 10.1101/2021.08.19.456963

**Authors:** Tomas Fitzgerald, Ewan Birney

## Abstract

Copy number variation (CNV) has long been known to influence human traits having a rich history of research into common and rare genetic disease and although CNV is accepted as an important class of genomic variation, progress on copy number (CN) phenotype associations from Next Generation Sequencing data (NGS) has been limited, in part, due to the relative difficulty in CNV detection and an enrichment for large numbers of false positives. To date most successful CN genome wide association studies (CN-GWAS) have focused on using predictive measures of dosage intolerance or gene burden tests to gain sufficient power for detecting CN effects. Here we present a novel method for large scale CN analysis from NGS data generating robust CN estimates and allowing CN-GWAS to be performed genome wide in discovery mode. We provide a detailed analysis in the large scale UK BioBank resource and a specifically designed software package for deriving CN estimates from NGS data that are robust enough to be used for CN-GWAS. We use these methods to perform genome wide CN-GWAS analysis across 78 human traits discovering 862 genetic associations that are likely to contribute strongly to trait distributions based solely on their CN or by acting in concert with other genetic variation. Finally, we undertake an analysis comparing CNV and SNP association signals across the same traits and samples, defining specific CNV association classes based on whether they could be detected using standard SNP-GWAS in the UK Biobank.

## Main

Genome wide association studies (GWAS) are a well-established genetic technique having made thousands of robust associations between traits and sequence level genetic variation (1–7). Often these associations can have significant impacts on the understanding, and in some cases, the treatment of human disease (8–10). However, for the majority of common genetic diseases these associations only account for part of the heritable disease risk (11–13). In terms of total base pairs differences CNV accounts for the majority of genetic differences between any two genomes (14–18) and have long been known to cause large differences in human trait distributions (19–21), often with a strong impact on human health (22,23). This is highlighted best within the large body of research that has spent decades studying CNV in relation to rare genetic disease (24–26). Although it is widely accepted that CNV can contribute significantly to differences in human traits, to date, methods for large scale CNV to phenotype association studies, ie the equivalent of GWAS for CNVs, rather than SNPs have been hampered by a number of factors, including methodological difficulties (27), the availability of sufficiently large datasets and the ability to interpret complex rearrangements from sequencing data (28,29).

Although CNVs have been a major component of routine clinical medical genetics screening for over a decade the interpretation of individual CNV events remains challenging (30,31), with most clinical testing laboratories routinely finding potentially pathogenic CNVs in patients with intellectual disability, autism spectrum disorders, and/or multiple congenital anomalies (32–34). Furthermore, although CNV detection from sequence data in a clinical setting is in active development, most clinical CNVs are still discovered using specialized microarray platforms (35). Most CNVs with strong effects are rare and are often discovered as *de novo* mutations in patients across a range of genomic disorders (36,37). Although the role of *de novo* CNVs in rare genetic disorders is well characterised, other types of genetic models such as X-linked, recessive, mosaicism, imprinting, digenic or even non-coding CNVs are still to be fully explored. Furthermore, it has been observed that the overall burden of CNVs is higher in specific patient groups compared to controls (38,39) indicating potential combinatorial CNV effects (40). It is conceivable that specific combinations of CNVs, that when observed in isolation would often be ignored, may, by acting in concert, have a large potential to cause phenotypic differences due to factors such as dosage compensation, incomplete penetrance and polygenic effects (41,42). Finally, the impact of CNVs in rare diseases is likely to be large, whereas one should expect weaker effects of all variation in more common diseases, consistent with their polygenic behaviour.

A large number of CNV specific genotype-phenotype correlations have been observed in relatively small-scale studies of specific patient groups (43) or by collaborative efforts to share genetic data for rare disease (44), however CNVs have also been associated with a number of complex diseases (45,46). More recently large scale CNV association testing from datasets such as the UK Biobank have found a number of highly significant loci in relation to certain human traits (47) and previous studies focused on cognitive traits such as schizophrenia (48) and autism (49) have demonstrated the utility of leveraging single-nucleotide polymorphism (SNP) genotyping arrays to search for novel CNV associations. Focussed studies into specific human traits have leveraged large scale SNP genotypes to perform association testing with great success (50–54) however these studies have often focussed on predefined lists of CNV regions known to be important within a clinical setting (53). Another important consideration is that SNP genotyping arrays in general have a limited resolution to detect small CNVs and a limited sensitivity for CNV discovery genome wide due to the distribution of SNPs across the genome and a limited dose response (55). Recently Auwerx et al showed both the power and limitations of genotype based CNV associations in the UK BioBank using the full 500,000 genotyping cohort, finding 131 CNV association signals across 47 phenotypes but with the smallest CNV detected of 49 kb at 1p36.11 that they found to be in association with reticulocyte count, platelet count and HbA1c (56). However, most of the CNV association signals detected involved large recurrent CNVs with a mean size of 817 kb again highlighting the known limitation in resolution when using SNP genotyping arrays for CNV discovery.

It is reasonable to assume that CNVs may account for a substantial portion of the variance observed in common disease risk. Some of these CNVs will be in strong linkage disequilibrium (LD) with SNPs, and so they can be discovered by tagging polymorphisms, but the causal change is impossible to narrow down only using SNPs. Other CNVs might not have good “tagging” SNPs and furthermore, recurrent CNVs are far more common than recurrent SNPs, with the CNV mutation rate estimated between 1.7 × 10⁻⁶ to 1.0 × 10⁻⁴ compared to between 1.8–2.5 × 10⁻⁸ per base pair per generation for point mutations (19,57), meaning that the aggregate higher frequency CNVs with the same functional impact are hard to model using the combination of rare haplotypes. With the advance of large data cohorts such as the UK biobank with vast amounts of sequence data sets that are amenable to copy number estimation (58–60) the ability to perform high resolution genome wide GWAS testing for CNVs has become more feasible. One challenge for large scale CNV discovery has been variability in raw sequencing depth due to other factors, most likely extraction techniques and immune system state at the time of blood draw. This variation gives rise to complex noise characteristics in raw sequencing read depth across the genome between samples, so called “genomic waves”. To explore this one needs robust large scale normalisation strategies for CNVs from sequencing data, an appropriate discovery method for CNVs and a way to easily integrate both CNV and SNP based associations into one framework.

In this work we address some of these issues by providing a new discovery method for CNVs, CNest, based on novel normalisation techniques for large scale cohorts. Rather than trying to create individual models of alleles for each CNV locus, we have chosen to use a straightforward linear model for discovery. This linear model is both consistent across all CNV loci and has many similar properties to the linear models used in SNP GWAS. As such we can use the same covariates, same diagnostic style QQ plots and place SNPs and CNVs into the same framework. Post discovery we show we can provide more detailed modelling of at least some loci. We provide a comprehensive CNV analysis using this method on the large UK biobank cohort with exome sequences. To explore the relationship with established SNP polymorphisms, we also performed both CNV and SNP GWAS within a single framework, applying our methods across exactly the same set of UK Biobank samples and then interrogating the resulting associations across a diverse set of traits.

We find many CNV to phenotype associations, though as expected many of these associations are also tagged by SNP polymorphisms. However, we have a subset of CNVs associations which cannot be discovered via SNPs and another subset which are coincident with strong SNP polymorphisms at locus, but not well correlated with any specific SNP. Finally, there are a large number of cases where the CNV is taggable, but the tagged SNP is at some distance from the CNV locus. Many of these associations recapitulate multiple known associations based on previous studies on both CNV and SNP genome association testing, whereas others discover new CNV specific findings in relation to the genetics of common human traits. We have made the software, CNest, which performs this discovery open source and provide portable workflows to run CNest, compatible with GA4GH standards (61).

## Results

### Copy Number Variation in 200,629 individuals from the UK Biobank

To identify exon resolution CNV regions across a large population of individuals from NGS data in the UK Biobank we developed a suite of flexible, highly scalable CNV analysis tools known as - **CNest**(**Methods**). Within this package we include a robust CNV caller as well as a set of tools and novel approaches to CNV association testing genome wide in discovery mode. A central component of these methods which is of critical importance for any accurate CNV detection, is the selection of appropriate reference datasets and normalisation procedures by modelling specific noise characteristics of WES/WGS, for example the presence and scale of genomic waves, to generate optimised relative copy number measurements across very large numbers of samples (**Methods**). After calling CNVs across the cohort of 200,629 samples with WES data we applied several important quality control measures to ensure that both the copy number measurements and CNV calls were consistent across this large cohort.

A subset of the diagnostic plots of CNest is shown in Figure 1. An obvious but important step in CNV analysis is the classification of sex based on the estimated copy number of the X chromosome. Well controlled 1 vs 2 copy number of the X chromosome indicates that the genomic wave control has worked successfully at least for the X chromosome (**Figure 1A**). A side effect of this analysis is the ability to detect sex chromosome aneuploidy. We detected 50 samples showing an unusually high number of copies on chromosome X (**Figure 1A**). These samples were assumed to be a mixture of data quality issues and real triple X cases. Triple X is a condition caused by random error during reproductive cell division and is found in approximately 1 in 1000 women. Although triple X has been associated with a number of trait differences, it can often go undiagnosed and depending on other social factors may never give rise to any noticeable problems (62). We also detected 51 datasets which show an unusual level of chromosome X coverage and can not be reliably assigned to either double (female) or single X (male). These we assume to be due to both inconsistent capture of chromosomal baits and potential mosaic sex chromosome events (ie. mosaic XXY). Sex chromosome aneuploidy is not a focus of this study and we simply exclude all samples that could not be reliably assigned to either double or single X based on their coverage profiles (**Figure 1A**); these copy number sex chromosome calls will be returned to the UK BioBank for further investigation by other investigators.

**Figure 1:**
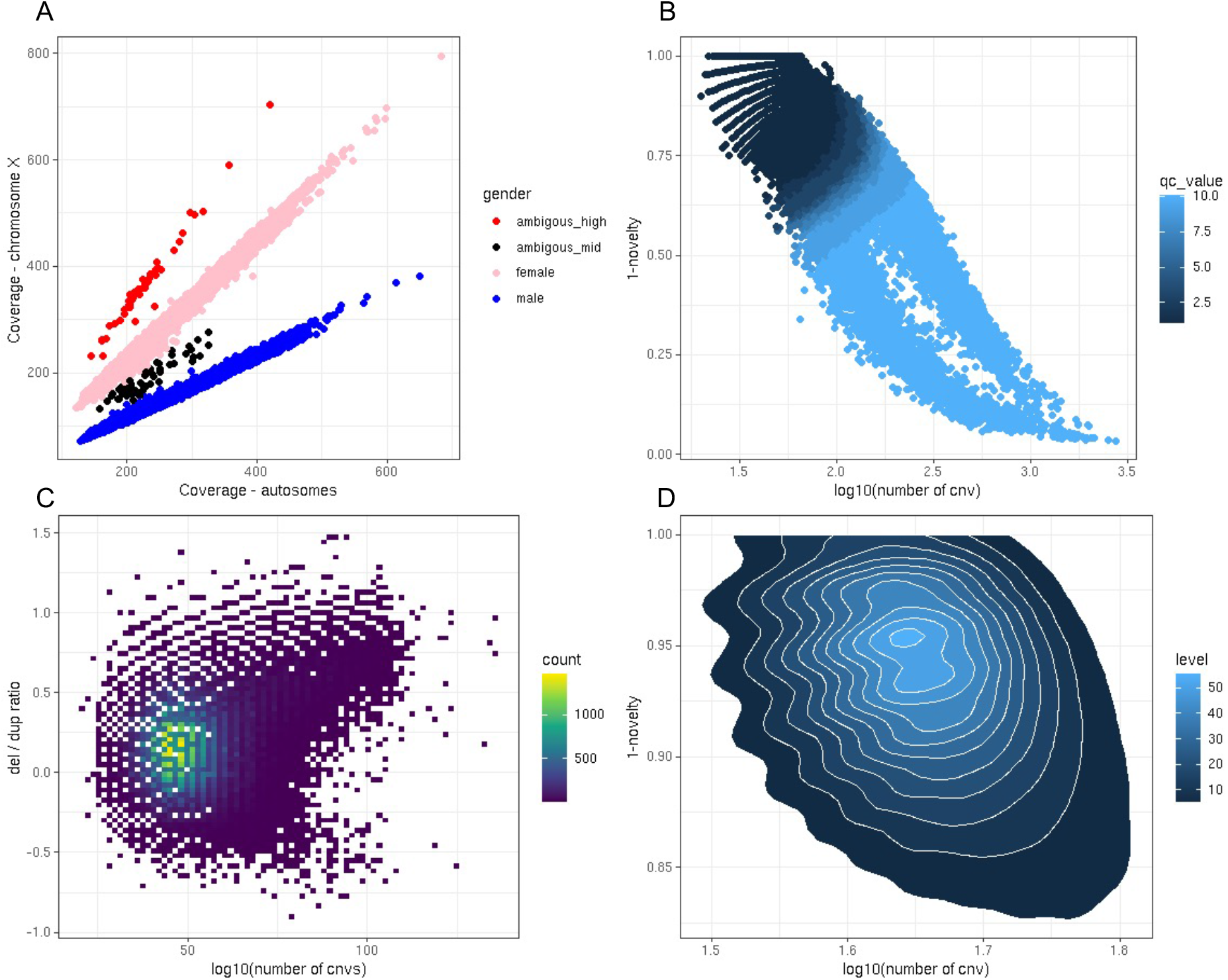
Quality control of CNV calls in the 200,629 UK Biobank exome sequences A: Gender classification, the relative coverage of autosomes compared to chromosome X across all samples. B: The total number of autosomal CNV calls vs a measure of the proportion of rare CNV calls per sample. C: The log10 of the loss to gain ratio vs. the log10 of the total number of CNV calls for each sample. D: A density plot showing panel B for all QC passed samples only.

Next some informative CNV quality information is contained within the consistency in the number of CNV calls in all samples against the proportion of those calls that are rare across the entire population (**Figure 1B**). This is similar to the genotyping extreme heterozygosity quality parameter used as standard in SNP genotyping QC. Given current estimates on the CNV mutation rate (63) we would expect very low numbers of *de novo* CNV events (less than 1 per genome) and rare CNVs to be infrequent in any individual genome, which is supported empirically here with a median of 3 rare CNVs per UK Biobank exome based on a 1% population frequency for loss and gains separately. For large scale CNV analysis in assumed healthy individuals it is sensible to assume that most genomes will on average display a consistent level of rare variation compared to the bulk of the population. Encouragingly after applying our strictest definition of quality control across greater than two hundred thousand exome sequences for CNV calling we obtain a greater than 92% pass rate indicating that for the vast majority of samples our CNV estimation and calling approach is highly consistent. When assessing these distributions in samples that passed our QC criteria the bulk of the data is tightly centred around a mean number of calls of 48 and a mean rarity rate of 0.07 (**Figure 1D**). Finally, there is no reason to expect, given known CNV formation mechanism such as non-allelic homologous recombination (NAHR) and non-homologous end-joining (NHEJ), that there would be any bias between the number of losses and gains when comparing large numbers of genomes in aggregate and although there are some outliers we observe a tight loss to gain ratio distribution with a median of 1.4 (**Figure 1C**).

To further assess some characteristics of the CNV calls we looked at how many predicted loss of function CNV (either deletion or truncating duplications) overlapped clinically important genes from the dd gene to phenotype (DDG2P) resource (24); we expect the common CNVs discovered in UK BioBank to be depleted in overlaps to these genes. Using a 50% reciprocal overlap rule against 218 mono-allelic loss of function genes from the DDG2P we found a total of 342 individuals CNV calls (**Supplementary Figure 1**), 40% of which were in the same gene *GLMN* which is known to be involved in Glomuvenous malformations (64). Overall, similar to previous work on pathogenic CNVs in the UK Biobank (52), we detect small numbers of CNVs in clinically important disease genes across the UK Biobank and rare variant analysis is not a focus of this study however we encourage interested researchers to make use of these high resolution CNV calls (**Data Availability**) where it might be possible to look at modifier effects for rare CNV events.

### Copy Number Variation Associations Testing in the UK Biobank

For CNV association testing genome wide in discovery mode we made use of both the copy number estimates and CNV calls generated by CNest and applied standard linear and logistic regression models using the copy number estimates as CNV dosage (**Methods**), analogous to the common dosage model of alleles from SNPs. Although the choice of linear models restricts our signal to sites displaying a linear relationship between copy number and trait, more sophisticated models that could have non-linear impacts on phenotypes can be complex to select and even more complex to analyse the resulting statistics consistently genomewide. Furthermore, this simple model is similar to those most often used in SNP GWAS (65) and so more easily jointly integrated with SNP discovery. All models were applied to unrelated samples from the PCA-defined European cluster (SNP PCs 1 and 2) and include standard covariates with 10 principle components (PCs) derived from both SNP and CNV estimates independently.

We performed CNV association testing genome-wide in discovery mode across 46 different main UK Biobank fields, including 30 quantitative and 16 binary traits across a variety of physiological, lifestyle and health related categories (**Supplementary Table 1**). We used diagnostic QQ plots and the associated genomic inflation statistic to be confident that our model produced a well behaved statistical test in which the majority of the genome fits the expected null hypothesis (**Supplementary Table 1**). In total, after fine mapping to select the most associated probe for each CNV-phenotype association at a locus (**Methods**) we discovered 646 significant CNV specific associations across 34 traits, 24 quantitative and 10 binary (**Supplementary Figure 2**). Additionally, we selected all instances of the “First Occurrences” UK Biobank field that had greater than 500 cases mapping to an ICD10 code (UK Biobank field 1712), resulting in 398 different codes that we then used as case/control labels for CNV association testing using our standard logistic regression models. These 398 labels covered 15 broader categories (**Supplementary Table 2**) and we obtained significant associations for 44 ICD10 codes across 13 of the broad categories with no significant associations for any of the traits under the categories, “Ear and mastoid process disorders” (field 2408) and “Congenital disruptions and chromosomal abnormalities” (field 2417).

Overall, across most UK Biobank traits that we tested which had sufficient numbers of samples we get strong discovery signals for CNV associations genome wide (**Supplementary Table 1**). When performing association tests on some well known traits we obtain significant, highly specific associations (**Figure 2**). Importantly all association signals that we obtained were by definition within exons and can be directly linked to genes, providing strong evidence for trait to gene mapping which has potential not only for discovering novel CNV association but also to provide strong supporting evidence for previous SNP GWAS results.

**Figure 2:**
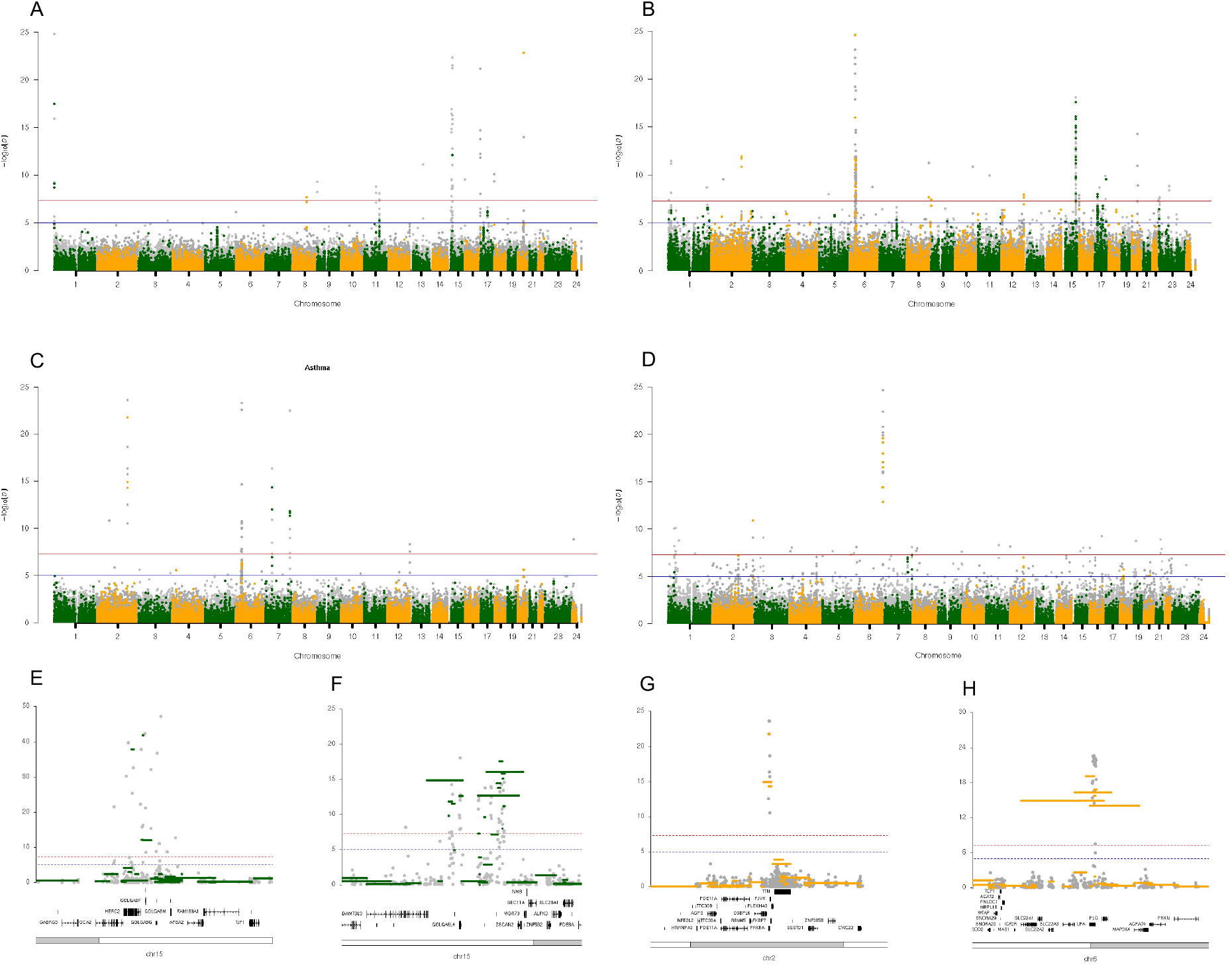
Copy number association Manhattan plots for four different UK Biobank traits, exon level signals are shown in different shades of grey and CNV calls signals in orange and green. A: Associations for hair colour using a linear model, B: Associations for standing height using a linear model, C: Associations for disease coding asthma using a logistic model, D: Associations for disease coding myocardial infarction using a logistic model. E: Zoom locus plot showing chr15 around the OCA2 / HERC2 genes for hair colour signal. F: Zoom locus plot showing chr15 around the ADAMTSL3 / UBE2Q2L / GOLGA6L4 genes for standing height signal. G: Zoom locus plot showing chr2 around the genes CHROMR, PRKRA and PJVK for asthma signal. H: Zoom locus plot showing chr6 around the LPA gene for myocardial infarction signal.

For example, for hair colour (**Figure 2A**) after fine mapping we detect 30 copy number variable regions that pass genome wide significance, the majority of which (20/30) have been found to be associated with pigmentation by previous SNP GWAS (**Supplementary Table 3**). A particularly strong signal is found in a region containing the *OCA2* and *HERC2* genes with a - log10 P > 230. Both OCA2 and HERC2 are well known genes involved in pigmentation in both humans and mice and have been shown to be involved in iris, skin and hair pigmentation in human (66–69). Pathogenic variation in *OCA2* including CNVs are known to be a leading cause of the rare genetic disorder oculocutaneous albinism and an increased susceptibility to melanoma (70–72). The strongest signal and lead exon (**Figure 1E**) around the fine mapped CNV region is found in the hect domain and RCC1-like domain 2 (*HERC2*) which has been well described by multiple studies and has a strong association to pigmentation of the eyes, hair and skin (66,73,74). Some of the novel associations we discover in relation to hair colour (**Supplementary Table 3**) include signals around 15q11.2, including several of the *GOLGA* genes which have functions relating to membrane traffic and Golgi structure; however, the precise function is unclear. Human chromosome 15 contains multiple copies of the GOLGA core elements close to the evolutionary conserved chromosome 15 low copy repeat (LCR15) duplicons (75) in primates at which several structural rearrangements break points have been described and linked to disorders and structural abnormalities such as Prader-Willi and Angelman syndromes (76). Furthermore, downstream *GOLGA* genes also detected here (*GOLGA8F*, *GOLGA8G* and *GOLGA8M*) have been linked to hair colour in previous SNP GWAS studies and from analysis of the UK Biobank (77).

Another example is standing height. After fine mapping our CNV signals into regions associated with standing height we discovered 45 distinct regions encompassing between 1 and 2 genes (**Figure 2B**). Again, the majority of these regions (27/45) contained at least one gene that had previously been associated with height from SNP GWAS (**Supplementary Table 4**) however with little to no evidence of CNV associations. The strongest signal outside of the HLA was seen in a region downstream of *ADAMTSL3* at 15q25.2, including the *UBE2Q2L* and *GOLGA6L4* genes and which is enriched for segmental duplications (78). There is a single exon signal at exon 15 of the well known *ADAMTSL3* gene; *ADAMTSL3* has previous evidence from multiple SNP GWAS studies of being associated with height (79–82) and with certain neurological disorders such as schizophrenia (83) and bipolar disorders (84) acting under a proposed alternative splicing mechanism (85). CNV at *ADAMTSL3* has yet to be described in relation to human height and interestingly this region contains multiple different CNV events varying in size all with strong association signals to height (**Figure 2F**). One for these CNVs overlaps exons 28-30 of *ADAMTSL3* which would be likely to result in truncation of the PLAC (protease and lacunin) domain (86). Strong human height CNV association signals are observed in the segmental duplication rich region downstream of *ADAMTSL3* including exons within the *UBE2Q2L* and *GOLGA6L4* genes both of which have been linked to neurological disorders by previous work (87,88) with *UBE2Q2L* also having been specifically linked with human height (81,89). Recurrent deletions and duplications at 15q25.2 have been described in relation to rare disease including neurological traits (90) however have not yet been described as a hotspot for structural rearrangements associated with common human traits such as height. We also found additional novel regions with no evidence of prior association to height (**Supplementary Table 4**), including one region at the Neuroblastoma BreakPoint Family *NBPF1* gene involving the highly copy number variable *DUF1220* domain (91) which have be previously associated in a dose-dependent manner with important human traits such as microcephaly and macrocephaly, brain size and neurological disorders (19,92,93).

Next we applied logistic regression models (**Methods**) to perform trait association testing using binary disease codes, again analogous to the equivalent SNP allele dosage test. First, we used the algorithmically defined health-related outcomes from the UK Biobank to perform case / control type association analysis for CNVs (**Figure 2 C and D**). For Asthma we discover 18 fine mapped CNV regions (**Figure 2C**). Strong CNV association signal was found around a region containing the 3 genes *CHROMR, PRKRA* and *PJVK* and upstream of *TTN* (**Figure 2G**). None of the 3 genes have previous evidence of a link to asthma however both *PRKRA* and *PJVK* have been found to be associated with lung functions such as vital capacity and forced expiratory volume (FEV) in a recent study (94). In contrast the cholesterol induced regulator of metabolism RNA *CHROMR* has no previous association to asthma and its precise function is poorly understood. Unsurprisingly the strongest signal for asthma is found in the HLA with the lead signal specifically restricted to the *HLA-DQA2* gene which has strong prior associations to asthma and hay fever all based on intergenic SNPs from multiple SNP based GWAS investigations (95–99). Some of our novel CNV associations for asthma (**Supplementary Table 5**) include signals in genes, *TAP2* and *STARD3NL*, which have not been linked with asthma by previous SNP GWAS studies but have evidence of association to certain other respiratory diseases (100,101).

For acute myocardial infarction (MI) we discover 26 fine mapped associations (**Figure 2D**). The strongest signal for MI is found within the *LPA* gene and this signal is found consistently across most heart related traits that we have tested in the UK Biobank. The *LPA* gene encodes a substantial portion of lipoprotein(a) and has been linked to numerous heart related diseases including coronary artery disease (CAD), aortic atherosclerosis and MI (102–104). Changes in dosage of the *LPA* gene, specifically the KIV-2 copy number alteration, has been previously linked to changes in lipoprotein(a) levels and a modified risk of heart disease (CAD) (105–108). This analysis of CNV association across a large cohort provides important additional information and allows a detailed estimate of the effect size for differences in *LPA* copy number in relation to the risk of MI. Interestingly we detect multiple different sized CNV events that hit the *LPA* gene (**Figure 2G**) but also include other coding regions with the lead exonic signal always restricted solely to the *LPA* gene. To our knowledge, only 2/26 fine mapped CNV associations (*LPA* and *BMP1*) have a direct association to MI from previous SNP GWAS testing (103,109) however a large fraction of the remaining regions have prior associations to other important heart related traits or cardiac disease risk factors (**Supplementary Table 6**). For example, the *TM2D1* gene that has prior association to electrocardiography (110) and the structure of the left cardiac ventricle (111); the *DPP6* gene that has been associated with multiple heart related phenotypes including sudden cardiac arrest (112); and genes associated with blood lipid level measurements such as *LCAT* and *RCAN1* (113,114).

For most of the UK Biobank main traits tested we discovered new CNV specific associations (**Supplementary Table 7**), for example, for the eye related trait corneal hysteresis we detect robust CNV associations in exons 15 to 36 of the *ANAPC1* gene (**Supplementary Figure 3**) in which sequence variation has been estimated to account for 24% of corneal endothelial cell density variability (115); and for both corneal hysteresis and intraocular pressure we discover exon level associations in the important *TCF4* gene which is known to be involved in several eye disorders such as fuchs corneal dystrophy (116). For red blood cell related traits we detect a large number of associations that have prior evidence of association from SNP GWAS (**Supplementary Figure 4**) such as variation in and around the *ABO* gene (117); for lifestyle measures such as alcohol consumption we find associations within known genes (118) such as *NPIPB6*; and for cognitive measures we also discover CNV association in genes with previous evidence of association from SNP GWAS testing in the UK Biobank, such as the *ARL17B* gene in association with reaction speed (119). All these CNV discoveries deserve integration with SNP polymorphisms and the often well studied biology around these loci and as described in the **Data Availability** section we have made all these results available to the community in a variety of ways.

### First Occurrences ICD10 Code Copy Number Variation Associations

To complement the UK BioBank measured and binary traits we also explore CNV associations to direct healthcare measures, as represented by the Hospital Episode Statistics (HES) captured data on ICD10 codes in the UK BioBank. We performed CNV association testing genomewide using the health related outcomes “First Occurrences” fields as case control labels for all codes that had greater than 500 cases within the UK Biobank 200K exome release. Although a relatively large number of those codes related to events that would be unlikely to show a genetic association, for example “other headache syndrome” and “bacterial infection of unspecified site”, we did not preselect or filter out any case labels and ran CNV association testing across all labels resulting in 398 cases control genome wide CNV association tests (**Supplementary Table 2 and 8**). Overall, across all 398 codes we discovered 242 CNV specific associations across 44 different codes covering 144 unique genes (**Figure 3A**). A large fraction of these associations (117/242) were located within the HLA super locus at 6q21 between chromosome positions chr6:30500001-46200000, and there were 6 traits that had no associations outside of the HLA super locus, 13 traits that had associations both within and outside of the HLA and 25 traits that had associations exclusively outside of the HLA.

**Figure 3:**
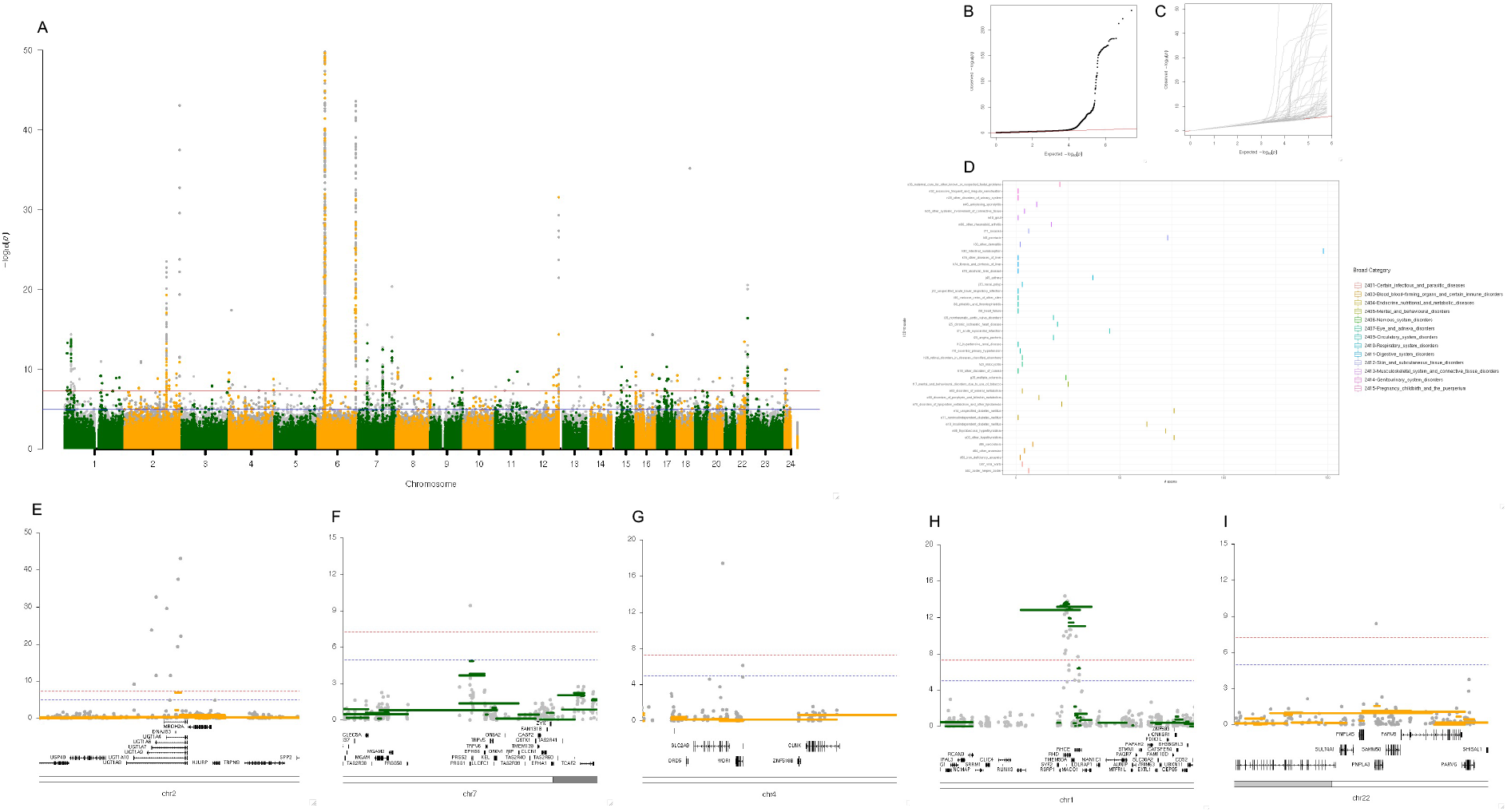
ICD10 code case/control copy number associations. A: Combined and overlaid manhattan plot for CNV associations across 44 ICD10 codes. B: Combined QQ plot including all p-values from association results across all 44 traits. C: Overlaid QQ plots showing all individual QQ plots for the 44 traits. D: Boxplots showing the number of significantly associating exons for all 44 trait association tests. E: Locus zoom plot at UGT1A genes for ICD10 code E80 - disorders of porphyrin and bilirubin metabolism. F: Locus zoom plot at the PRSS1 gene for ICD10 code D50 - iron deficiency anaemia. G: Locus zoom plot at the SLC2A9 gene for ICD10 code M10. - Gout H: Locus zoom plot at the RHD and RHCE genes for ICD10 code O36 - maternal care for known or suspected foetal problems. I: Locus zoom plot at the PNPLA3 gene for ICD10 code K74 - fibrosis and cirrhosis of liver.

After fine mapping the majority of ICD10 codes (27/44) had between 1 and 2 significantly associating regions with 9 ICD10 codes having between 3 and 10 and 8 ICD10 codes having greater than 10 associations. Most fine mapped regions were small (**Supplementary Table 8)** with a median number of significant exons of 3 per fine mapped region (**Figure 3D**) with the largest region involving 52 exons across 5 different genes in association with ICD10 code K90: intestinal malabsorption. Almost all the association results were very well controlled with inflation factors (lambda) ranging from 0.984 to 1.140 except for ICD10 code F17: mental and behavioural disorders due to use of tobacco which showed mild inflation with a lambda of 1.382 (**Figure 3 B, C and Supplementary Table 1**). In total we detected 242 associations ranging from well known important regions of the genome, through to completely novel findings based on CNVs alone. All association results and summary manhattan plots for the 44 significantly associating ICD10 codes are provided in **Supplementary Material**.

For example, for ICD10 code E80: disorders of porphyrin and bilirubin metabolism, we found multiple strong signals involving specific exons across several UDP-glucuronosyltransferase genes (*UGT1A10*, 9, 8, 7, 6 and 4) (**Figure 3E**). Genetic variation of *UGT1A* genes has been associated with disorders of bilirubin metabolism including Gilbert’s syndrome by multiple previous SNP GWAS studies (120,121) with, for example, very strong association signal at *UGT1A10* for the intron variant rs6742078 (2_233763993_G_T) (122). This specific SNP has also been linked to other lipid metabolism disorders such as Gallstones Disease (GSD) (123) and although studies looking at CNV burden analysis of lipid metabolism genes has shown a significant enrichment in GSD cases none of those associations could be attributed to any single gene (124). Here we provide novel CNV associations at *UGT1A* genes with a direct link to bilirubin metabolism that could be an important risk factor for lipid metabolism related disorders.

Another example is ICD10 code D50: iron deficiency anaemia we discovered two significantly associated loci on chromosome 7 (**Figure 3F**) one of which covers exons 4-6 of the cationic trypsinogene gene *PRSS1* that has been linked to chronic pancreatitis by multiple studies (125,126). Autosomal dominant mutations in *PRSS1* are thought to be a leading cause of hereditary pancreatitis, a rare condition that results in recurrent inflammation of the pancreas, and an increased risk of pancreatic cancer (127). As such *PRSS1* is regularly tested in patients with suspected hereditary pancreatitis (128) however the *PRSS1* gene contains multiple known variants, including copy number changes, often with unknown clinical importance (129). Iron metabolism and pancreatic function are closely related processes (130) with evidence that pancreatic enzyme levels influence the efficiency of iron absorption (131). Here we provide a link between the copy number at exons 4-6 of the *PRSS1* gene with the ICD10 code D50 relating to iron deficiency anaemia that may be a result of pancreatic dysfunction.

For Musculoskeletal disorders we discovered 17 fine mapped association loci across 4 different traits including one location at 4p16.1 at exon 3 of the *SLC2A9* gene that was associated with ICD10 code M10: gout (**Figure 3G**). Gout is a swelling of joints, normally in the feet, that is caused by hyperuricemia (an excess of uric acid in the blood) with mutations at *SLC2A9* having been found to be associated with serum urate concentrations and the onset of gout (132,133). A non-coding CNV near *SLC2A9* (integenic and approximately 200 kb upstream of the *SLC2A9* gene) has been described in association with serum uric acid levels (134) however CNVs in coding regions of the *SLC2A9* have not yet been discovered in relation to uric acid level or with a direct association to gout. Here we provide a novel CNV association result at exon 3 of the *SLC2A9* gene with a direct association to gout from the UK Biobank.

For Pregnancy childbirth and the puerperium, we discovered fine mapped CNV associations against code O36: maternal care for known or suspected foetal problems at 1p36.11 including the *RHD* and *RHCE* genes (**Figure 3H**). Variation and *RHD* gene deletion in the human Rh blood group system has been extensively studied in relation to pregnancy risk (135) where prior to the development of medical treatments, Rh-negative (D-negative) mothers were at significant risk of haemolytic disease of the newborn (HDN). It is still unclear what potential benefit the RHD gene deletion may have that merits its relatively high frequency in the human population (136). Blood tests are normally carried out in D-negative expectant mothers to determine the Rh factor status of the child and direct treatment if using anti-D injection is required (137). However variation in the less well understood Rh C and E alleles of *RHCE* is clinically relevant, influences the risk of HDN (138) and this association discovered in this study merits further investigation for this well understood risk factor for pregnancy.

For ICD10 code K74: fibrosis and cirrhosis of liver we discovered a single CNV association at exon 3 of the *PNPLA3* gene (**Figure 3I**). Cirrhosis of the liver is a disorder in which the liver parenchyma is replaced with fibrous tissue and is often caused by alcoholism as well as hepatitis B and C infection (139,140). The *PNPLA3* gene has been found to be associated with liver cirrhosis by multiple SNP GWAS studies (141,142) and although CNVs at *PNPLA3* has not been well described or linked to Cirrhosis in the past it has been shown that transcriptional regulation of *PNPLA3* has an impact on liver disease with higher levels of *PNPLA3* mRNA in the cytoplasm being negatively associated with the severity of alcoholic fatty liver disease (NAFLD) in humans (143).

Finally, we looked across 5 additional heart related ICD10 codes (I20, I21, I25, I35 and I50) and found strong CNV association signals at the *LPA* gene (see above) for all cases with the exception of I50: heart failure (**Supplementary Figure 5**). When comparing signal strength between the 5 heart related ICD10 codes we observe a clear sample size effect with the tests showing the stronger signals tending to have larger number of cases (**Supplementary Figure 5 B-E**). The five ICD10 codes included were I25: chronic ischaemic heart disease (20,503 cases), I20: angina pectoris (10,117 cases), I21: acute myocardial infarction (3,698 cases), I35: nonheumatic aortic valve disorders (1,692 cases) and I50: heart failure (3,557 cases). CNV association at LPA is a major feature in all heart related phenotypes we have tested in the UK Biobank providing further evidence that changes in dosage of *LPA* is a significant risk factor for heart disease in humans.

In summary we discovered 862 new fine mapped CNV associations across 78 different traits (24 quantitative and 54 binary) using the 200K whole exome release from the UK Biobank the majority of which have either been previously discovered by SNP GWAS tests or have compelling evidence from other research areas such as health care, rare disease or animal models (**Supplementary Material**), but a significant minority are entirely novel. These new association results and genome wide association testing approach for CNV provides important new insights into the contribution of CNV in complex human traits which in some cases can have a direct relevance to health related outcomes and genetic risk profiles. We encourage interested readers to pursue the discoveries discussed here and listed in our supplementary information further.

### Combined CNV and SNP based associations in the UK Biobank

To investigate the relationship between SNP and CNV associations in the UK Biobank we ran standard SNP based GWAS tests across 6 quantitative traits using the same sample set as we used for CNV association testing (**Methods**). The intention is to allow a direct comparison of SNP to CNV association signals across a variety of human traits. This allowed us to start to explore the underlying genome landscape for associations and to classify individual associations into those that were detectable by SNP and CNV GWAS independently against those that are specific to CNVs. We set out to classify all fine mapped CNV association regions using the association results from SNP GWAS tests in the same samples and traits as well as calculating the r^2 between the lead exon CNV signal and all SNP genotypes within 1 MB. To do this we applied a simple set of rules considering signal strength, correlation between SNP and CNV genotypes and distance (**Methods**). Using this approach we classified CNV signals into *CNV only* - signals that were detectable by CNV GWAS only, *CNV allele* - signals that were present at the same locus by both SNP and CNV GWAS but with very little correlation between them, *SNP-CNV near* - signals that were detectable by both SNP and CNV GWAS and where those signals were highly likely to be assigned to the same gene and *SNP-CNV far* - signals that could be detected by both SNP and CNV GWAS but were highly likely to be assigned to different genes.

Across a total of 133 fine mapped CNV association regions we classified 17% (23/113) as *CNV-only,* 44% (59/133) as *CNV-allele*, 28% (38/133) as *SNP-CNV near* and 11% (13/133) as *SNP-CNV far* (**Supplementary Table 9**). It is worth noting that we choose to set the r^2 cut-off relatively low since very strong SNP-CNV tagging is rare genome wide and we preferred to be strict in the definition of novel CNV events (*CNV-only* and *CNV-allele*) such that when we define an association as *CNV-allele* we can be sure that it is not well tagged by any SNP that associates with the same trait and that for *CNV-only* types there is no SNP based association that passes genome wide significance within 1MB around the lead CNV position. We consider the *SNP-CNV far, CNV-allele* and *CNV-only* association type as different types of novel CNV associations whereas for the *SNP-CNV near* type we assume that the signals from both variant types are highly likely to be tagging the same functional variant; nevertheless, the CNV association may well provide functional insight for the locus.

To present the relationship between CNV and SNP based association results we created a similar display to the locus zoom plots from SNP GWAS but with the focal point being the lead exon from the fine mapped CNV region. We show 1 example (**Figure 4**) and an additional 2 examples (**Supplementary Figure 6**) for each of the CNV association type classifications covering a variety of SNP-CNV correlation patterns and differential signal strengths. A general feature in all cases is that all CNV association signals are in coding regions whereas SNP association signals are most often observed in non-coding regions, either intronic or intergenic.

**Figure 4:**
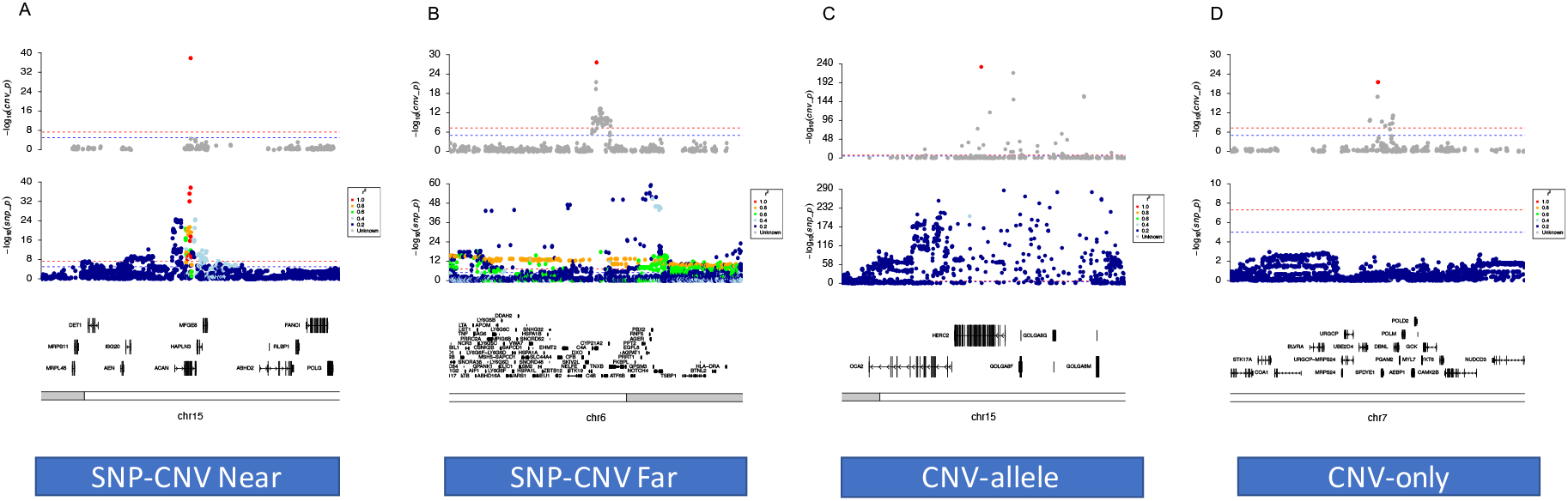
Locus zoom plots showing SNP and CNV association results for the different CNV association type classifications for quantitative traits. A: SNP-CNV near association plot for standing height at ACAN. B: SNP-CNV far association plot for FEV/FEC ratio at C4A. C: CNV-allele association plot for hair colour at HERC2. D: CNV-only association plot for chronotype at SPDYE1.

For standing height, we show clear complementary signals at the *ACAN* gene for SNP based and CNV based association results (**Figure 4A**), an example of the *SNP-CNV near* class with the CNV signal being well tagged by 30 SNPs within the gene body between exons 6 to 12. The CNV signal is restricted to exon 12 that encodes the chondroitin sulfate attachment (CS) domain (144) which is important for aggregation with hyaluronan resulting in a strong negative charge that gives rise to load-bearing properties of cartilage (145). Mutations in *ACAN* have been studied in relation to both syndromic and nonsyndromic human traits with a number of different impactful variants having been discovered (80,146,147). Earlier studies found *ACAN* to be a strong candidate for autosomal dominant disorders such as spondyloepiphyseal dysplasia Kimberley type (SEDK) and early-onset osteoarthritis (OA) from genetic linkage analysis and mouse models of chondrodysplasia (144) however, heterozygous mutations in *ACAN* display highly variable nonsyndromic phenotypes including short stature, early onset osteoarthritis and mild dysmorphic features (148). In this case, although the CNV is both well tagged and close to the SNP associations, the CNV directly suggests the functional variant of these SNPs is the deletion of this exon implying that haploinsufficiency of the *ACAN* gene is the main mechanism underlying these associations.

A *SNP-CNV far* example is shown in **Figure 4B**, being a 60KB region on chromosome 6 that is associated with the lung function measure FEV/FEC ratio including the *STK19, C4A, C4B* and *CYP21A2* genes with the lead exon CNV signal encompassing exons 26-30 of the *C4A* gene (**Figure 4B**). Interestingly, there are 208 tagging SNPs that pass genome wide significance for FEV/FEC ratio however none of these SNPs are located closest to either of the *C4* genes (*C4A* or *C4B).* The C4 genes encode an important part of the immune complement system and deficiencies (including CNV) at *C4* genes have been strongly associated with immune disorders such as Systemic Lupus Erythematosus (149,150). The *C4A* and *C4B* genes encode different components of the highly polymorphic C4 complement protein and can be distinguished from each other by four specific amino acids at positions 1101–1106 (151). Due to high sequence similarity the total copy number of *C4* can be defined as the sum between *C4A* and *C4B* (152), however both *C4A* and *C4B* are multiallelic CNV locations displaying common differences in copy number with *C4A* ranging between 0 to 5 and *C4B* between 0 to 4 copies (153). The CNVs at *C4* has not been linked previously with lung function; however, a recent study into chronic obstructive pulmonary disease (COPD) in the Korea Associated Resource cohort has investigated genome wide SNP interactions mapped to *C4B* in relation COPD and the FEV/FEC ratio measure (154). We provide new evidence for the role of *C4* CNV in lung function as measured by the FEV/FEC ratio in the UK Biobank.

Next, to investigate which gene to trait associations would be detectable by CNV association testing only, first we classified novel CNV associations as *CNV-allele* (detectable by both SNP and CNV GWAS but with no tagging) at the *OCA2/HERC2* locus, which is described above. Interestingly, although there is strong evidence of association, with similar signal strength, from both CNV and SNP based tests at *HERC2* there is very little correlation between the copy number estimates and SNP genotypes across the locus (**Figure 4C**) suggesting that these associations are likely to be operating via different functional mutations.

We also looked at events that would simply not have been detected by standard SNP GWAS tests in the same samples and classified these as *CNV-only*. An example is associations to chronotype. We discover a highly specific association within the *SPDYE1* gene on chromosome 7 where there are no tagging SNPs and no SNPs associations passing genome wide significance within 1MB (**Figure 4D)**. The *SPDYE1* gene has no previous association to measures of sleep patterns and very little description of CNVs at this location, however, a related gene *SPDYE6*, has been associated with insomnia via SNP based association testing in a much larger sample set (1.3M samples) (155). Furthermore, we discovered significant *CNV-only* associations at *SPDYE6* and the directly adjacent region containing the *POLR* and *SPDYE2B* genes which have been previously associated to chronotype by SNP GWAS tests in the UK Biobank (156). Here we provide strong evidence that CNVs at *SPDYE1, SPDYE6* and *POLR* related genes are associated with a measure of chronotype in the UK Biobank and may act in a dose dependent manner to influence an individual’s sleeping pattern.

We also present two additional examples of each of the different CNV association classes (**Supplementary Figure 6**), including; SNP-CNV near associations to hair colour at the *SPG7* gene (157) and to alcohol consumption at a 0.8MB region including 4 fine mapped CNV regions involving the *NPIPB6, NPIPB7, NPIPB9* and *SH2B1* genes (118,158–160); SNP-CNV far classifications for hair colour at the *TRIM49C* gene and standing height at a fine mapped region involving the *EVPLL* and *LGALS9C* genes; CNV-allele association types for heel bone density at the *WNT16* gene (161–164) and the FEV/FEC ratio at the *HTR4* gene (89,94,165–167); and CNV-only classifications for the FEV/FEC ratio at the *ZDHHC11B* gene (94) and for standing height at the *CDK11A* gene.

### Competitive SNP-CNV Association Models

To look in more detail at the relationship between SNP and CNV associations in the same samples and traits we performed some joint competitive association models for standing height and hair colour (**Methods**). Across 91 exon level CNV association signals that had at least one significant SNP within 1MB (and so would fall into the *CNV-allele*, *SNP-CNV Near* and *SNP-CNV Far* categories but excluding the *CNV-only*) around the lead CNV position, we performed eight different types of models (**Methods**). First, we applied a 3 component mixture model to the normalised copy number estimates (log2 ratio) to define copy number genotypes (assuming a simple deletion/gain process), analogous to the 3 SNP allele categories, relating to low (loss), medium (normal) and high (gain) copy number states. Next, we performed joint SNP and CNV competitive models using both the SNP with the highest signal strength against the same trait, in the same samples and the SNP showing the highest r^2 against the lead CNV exon within 1MB.

Overall, for the majority of signals (71/91) the denoising of copy number estimates into a 3 component copy number state model resulted in a slight lowering of signal with 17/91 sites dropping below genome wide significance (**Figure 5 A and B**).

**Figure 5:**
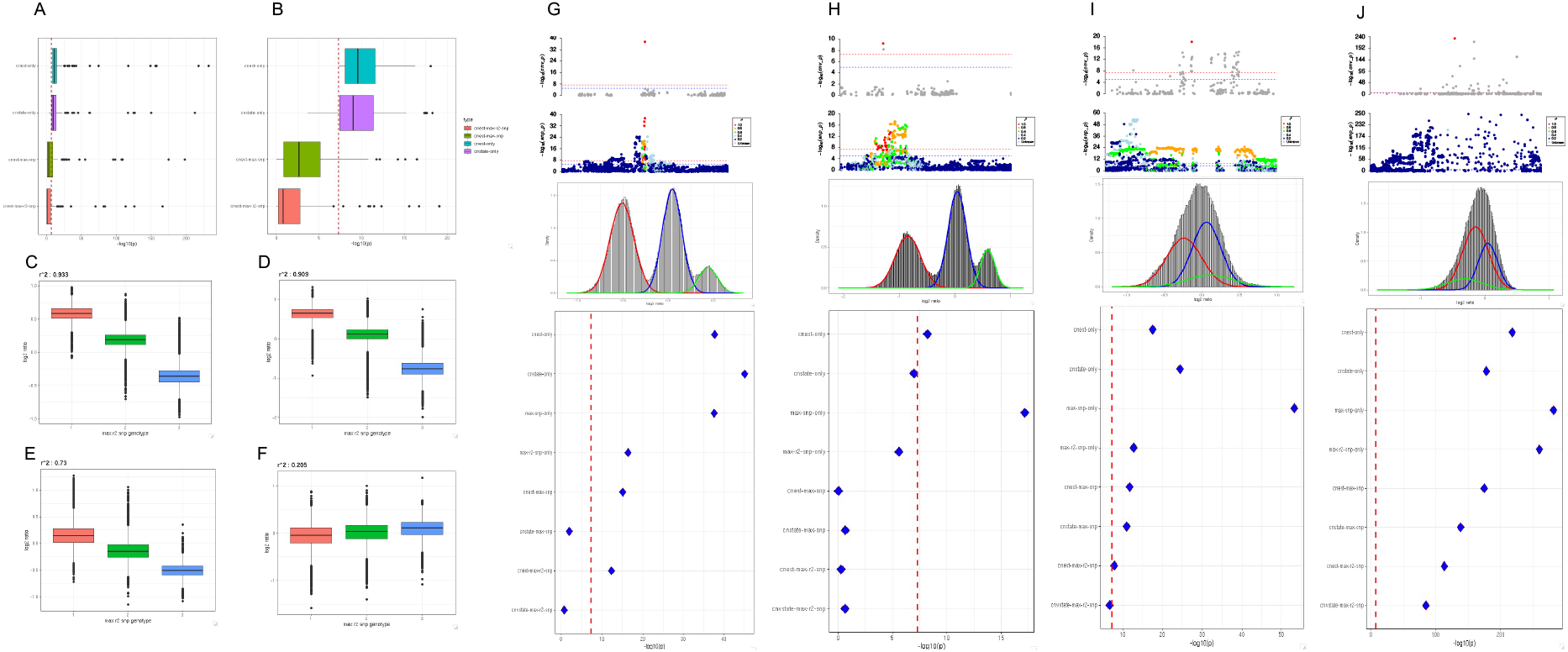
Competitive models for CNV and SNPs using copy number estimates, copy number genotypes and joint models including SNP genotypes from the most highly correlated SNP or the SNP with the highest association signal for the same trait within 1MB. A: Minus log10 p values for four different models, cnest-only: copy number estimates only, cnstate-only: copy number genotypes (3 component mixture model) only, cnest-max-snp: joint model with copy number estimates and the SNP with the highest association signal for the same trait within 1MB, cnest-max-r2-snp: joint model with copy number estimates and the the most highly correlated SNP within 1MB. B zoomed in view of panel A restricting the x-axis to a maximum −log10 p value of 20. Panels C - F: SNP genotypes from the most highly correlated SNP against the copy number estimate (log2 ratio) for 4 individual exon level association signals. further details of which are shown in panels G through to J. Panels G - J: Finer grain details for joint models of 4 exon level copy number association signals, top panel shows the copy number estimate association signal with the lead exon highlighted in red, second panel show the SNP genotypes association signal from SNP GWAS tests in the same samples and trait coloured by the r^2 of SNP genotypes against the lead exon signal from the CN GWAS, third panel shows the copy number estimate (log2 ratio) of the lead exon association from the CN GWAS fitted using a 3 component mixture model to define copy number genotypes, finally the fourth panel shows the −log10 p value from 8 different types of association model: cnstate-only, cnstate-only, max-snp-only, max-r2-snp-only, cnest-max-snp, cnstate-max-snp, cnest-max-r2-snp, cnstate-max-r2-snp.

For most sites the association signal strength was slightly lowered with a median −log10 p value reduction of 1.7 for the 3 state copy number model overall, however there were 20/91 sites that showed a marginal improvement is association signal strength with a median increase of 0.65 and one site at the *GOLGA6L4* gene showing the relatively large increase of a −log10 p value of 7 when using the copy number state compared to the copy number estimate model (**Figure 5I**). The drop in p values is to be expected due the selection of the most associated exon (winner’s curse phenomena) but it shows that there are not large gains to be made by a more categorical model for discovery.

Next, we performed some competitive SNP and CNV association models to further investigate the relationship between SNP and CNV associations across the 91 significant CNV locations. The simplest hypothesis is that sites that are well tagged and show association to the same trait will be most likely to be able to control each other’s signal in a pairwise competitive model. Across the 91 sites including the SNP with the highest signal strength irrespective of its r^2 can fully control 72% (66/91) of CNV associations resulting in the CNV signal being reduced below genome wide significance in the competitive models (**Figure 5 A and B**). Furthermore, when switching the SNP to those with the highest r^2 irrespective of association strength we observe an increased level of control with 80% (73/91) of CNV signals being reduced below genome wide significance. Indeed, the most frequent situation is that SNPs that can well tag the CNV estimates in aggregate (across all samples) are best able to control the CNV association signals within a competitive model with only 2 cases where highly correlated and significant SNPs are unable to fully control the CNV association (**Figure 5 G and I**). As expected for the 18 CNV associations that cannot be fully controlled by either SNP type the vast majority show very little tagging with 80% of these sites having a maximum r^2 below 0.6 for all SNPs within 1MB. However, there are two cases where the main assumption that highly correlated nearby SNPs will be able to control the CNV association does not hold true (**Figure 5 G and I**).

We show four examples of different types of association control from these competitive models, first a case where neither SNP type included in the models can fully correct the CNV estimate signal where we assume the SNPs are tagging the CNV (**Figure 5G**). Next an example where both SNPs, either the most significant or the most highly correlated, can fully control the CNV association signal and we assume that the CNV tags the more significant SNP (**Figure 5H**). It is worth noting that the most significant SNP in found in intron 1 of the *DOCK8* gene whereas the CNV signal and the tagging SNP are both found in the neighbouring *CBWD1* gene and that both genes have been found to be associated with the trait (hair colour) by previous SNP GWAS testing (74,157). We also show an example of a CNV association where neither SNP can fully control the copy number estimate signal, but the highest correlated SNP is able to push the copy number state joint model down just below genome wide significance (**Figure 5I**). The highest signal SNP is found closest to a different gene upstream of the CNV signal and there are multiple tagging SNPs surrounding the CNV locus which we assume all tag the CNV association. Finally, we show an example of a highly significant CNV association signal that has very little SNP tagging around the locus and is as expected not able to be controlled by either SNP in the competitive joint models supporting our general assumption that SNPs that cannot well tag nearby CNVs are unlikely to be able to control any CNV association even if both variant classes show significant associations to the same trait (**Figure 5J**).

Here we have shown results from combined SNP and CNV association testing across close to one hundred significant exon level CNV associations. We have shown that the most obvious assumption that nearby tagging SNPs are often able to control CNV associations holds true, but that more complex situations exist where aggregate variant correlations are not sufficient to predict the interactions between them in relation to trait association testing. We have also shown that it is possible to use copy number estimates in a dosage dependant linear model as a reasonable proxy for the underlying copy number state distribution and by applying similar methods to the highly successful SNP GWAS approach it is possible to discover novel CNV associations, add additional supporting evidence for SNP based trait association mapping and to further delineate the underlying genome architecture and variant interactions for trait associations.

## Discussion

In this paper we present a robust CNV to phenotype discovery process which is analogous to the traditional SNP based GWAS. A key foundation is a robust normalisation procedure which can handle the diversity of DNA presentation and extraction states in a large cohort. Armed with this normalised copy number level, we decided to model the complexity of CNVs in the genome as linear dosage variable present across the genome; this model is obviously an approximation to the reality of structural variation but allows consistency of statistical approach and the same degree of freedom across the genome, and means that similar covariates, methods and diagnostic procedures, with similar expected null model properties to SNP-GWAS can be used. Our resulting linear dosage model produces well calibrated statistics for both quantitative and qualitative traits, where the majority of associations fit the expected null model. The minority of associations where one can confidently reject the null model at a genome-wide significance level includes many well known individual CNV associations with a total of 862 associations across 78 different human traits. Using a subset of traits to perform CNV and SNP association testing in exactly the same samples, we found that 11% of these novel associations are tagged by SNPs though at some distance from the CNV association, 44% of associations are hard to discover using tagging variation, representing likely novel alleles at previously discovered loci, 17% of associations having no significant signal from any nearby SNP and the remaining 28% of associations are close to SNP polymorphisms which tag the CNV.

In this paper we have illustrated the large scale discovery of CNV associations with 12 examples in the main text and an additional 18 examples in the supplementary information. The examples vary from well established CNV loci (eg, *LPA* with heart disease loci) through partially understood CNV loci (eg the *UGT1A* gene in porphyrin and bilirubin metabolism) to very credible associations to paralogues with the same phenotype (the RCHE gene in pregnancy complications) or credible novel alleles in a gene with robust association to a phenotype (HERC gene with hair colour). There are numerous other examples in supplementary information, and the full information of the discovery processed here is available both via UK BioBank return of results and also via GWAS Catalogue. Even so, we have chosen only a subset of phenotypes present in UK BioBank, itself only one cohort; to enable broader discovery by others we have released CNest as an open source package and provided portable workflows consistent with GA4GH standards for other groups to use. We hope to build out user friendly tutorials, including the practical and necessary aspects of QC before discovery. We encourage the community to examine the results we have presented here and use the software to make more discoveries.

As well as performing large scale CNV specific association testing we were also able to investigate the correlation between SNP and CNV discoveries by performing SNP and CNV association testing independently and jointly using exactly the same sample sets and traits. Here it is possible to start to estimate the contribution that both types of variations have on trait associations in a large cohort and to gain some insight into the different types of interactions that can occur. As expected, for the CNV associations with some level of correlation to a SNP there are complex relationships between SNPs and CNVs; in some cases the SNP associations can completely “explain” the CNV association, whereas in other (rarer) situations the CNV association cannot be recapitulated by any particular SNP. This latter case is consistent with multiple CNVs arising on different haplotypes, where the CNV association appropriately aggregates the CNV information in a way which is far harder to achieve via tagging SNPs of each individual CNV. Even in the cases where the loci are discoverable by SNP methods, and the SNPs tag the CNV, the large expected impact of deletion or expansion of an exon makes CNV an interesting potential functional change.

The ability to jointly model SNPs and CNVs in the same framework will more easily allow for integration of these two types of variation. An obvious extension is to polygenic risk scores for traits, where the linear model for CNVs naturally fits with the additive linear SNP loci in a PRS. However, care needs to be taken over ascertainment and modelling for polygenic risk scores, in particular for certain traits such as blood based cancers where the normalisation procedures we have employed for CNV association might not be robust enough to distinguish germline from somatic changes in cancer risk. Importantly this means care needs to be taken about the time of blood sample compared to the onset of diseases in constructing such PRSs. Another extension will be using these linear variables as instrumental variables in Mendelian Randomisation techniques to understand causality between physiological processes and often disease outcomes. A similar concern on normalisation techniques needs to be considered, along with careful consideration of the assumptions behind any instrumental analysis.

Copy Number Variation has long been known as an important aspect of germline DNA variation and has long been a key part of rare genetic disease discovery and diagnosis. The system we have proposed here, CNest, can robustly find associations of CNVs to common phenotypes in large cohorts, but we have only started in providing a full catalog of these results. We encourage the community to explore the discoveries we have made in this paper, to use CNest to make more CNV associations in both UK BioBank and beyond, and to help extend the CNest method further to provide a more comprehensive view of human variation.

## Methods

### Sample Cohort and Phenotypes

For this study, we used 200,624 Whole Exome Sequencing datasets from the UK Biobank 200k release generated using the IDT xGen Exome Research Panel v1.0 including supplemental probes and sequenced with dual-indexed 75×75 bp paired-end reads on the Illumina NovaSeq 6000 platform using S2 and S4 flow cells (168). We used the aligned CRAM files from the OQFE pipeline which aligned and duplicate-marked all raw sequencing data (FASTQs) against the full GRCh38 reference in an alt-aware manner as described in the original FE manuscript (169). These aligned sequence datasets were used as the primary input in the CNest pipeline (**Details below**) for exome-wide copy number estimation and CNV calling. Phenotypes were extracted and linked to the copy number data under UK Biobank application number 49978, resulting in a total of 78 different traits (24 quantitative and 54 binary) that we tested for CNV association (**Supplementary Table 1**).

### Genetic data processing and copy number estimation

We used CNest (full source code available: https://github.com/tf2/CNest.git) to carry out large scale copy number estimation in the UK Biobank 200k WES release. This program was designed to provide accurate copy number estimation from very large NGS (WES and WGS) datasets. The first required step is to extract read coverage information for all genomic locations of interest, to do this CNest makes use of the samtools and htslib libraries (170) implementing a custom coverage extraction method that importantly filters reads based on several samtools alignment flags. The main flags of interest are BAM_FPROPER_PAIR, BAM_FDUP and BAM_FSECONDARY where we ensure that aligned reads have a MAPQ greater than 1, are primary alignements with proper pairs and are not PCR duplicates (**CNest docs**).

After extraction of coverage information, the first important step is to classify the sex of each sample based on the relative coverage on chromosome X. Here CNest implements a simple k-means clustering for the initial classification and quality control steps. This initial step results in the classification of two states relating to 2 or more and 1 or less copies of chromosome X (although less than 1 copy of chromosome X is biologically incompatible there can be data quality issues to account for when processing large volumes of data). CNest also implements a prototype automated classification to detect sex chromosome auniopoly which is based on the ‘abberant’ cluster (171) however we highly recommend that the sex clssification in checked by a human before moving onto the next steps. This is because all datasets are different and will contain a variety of sex classification types (**Figure 1A**), as standard across large numbers of samples we observe 3 types of sex chromosome dosage exclusion types and classify samples into either male or female, or ambiguous low, ambiguous mid, ambiguous high. All samples that are not classified as either male or female are removed from the subsequent steps.

The next crucial step is to derive sample specific dynamic reference datasets, briefly, CNest uses an optimisation process to select groups of appropriate datasets to make up individual references (or baseline) estimates across the genome for each sample individually. During these process CNest estimates the overall correlation of coverage information between all samples, applies a wavelet model for estimating the scale of genomic waves, and implements a dose response optimisation using sex mismatched samples and the expected single copy dosage change. This process has been designed to be extremely efficient across very large numbers of samples and results in a ranked list of which samples are most correlated in terms of certain coverage patterns and noise characteristics which are assumed to be the ideal set of samples to generate the baseline estimate. The only parameter needed to be decided on at this stage is the total number of datasets to use to generate the dynamic references, for the UK Biobank 200K release we elected to use 2,000 samples within each of the ~200K dynamic reference datasets. Although it may, in some cases, be preferable to allow the dynamic reference sets to be made up of different numbers of individual samples (which is possible using CNest) we decided to fix this number across all datasets as we wanted to minimise any potential biases within the resulting copy number estimates that could be introduced due to using differentially sized reference datasets.

Following the dynamic reference selection process, the median coverage for all genomic locations across all relevant reference datasets is calculated for each sample individually for matched, mismatched, and mixed sex classifications. These data values are stored in the custom CNest binary format to allow fast random access across the genome and across different sample sets. Finally, the coverage information and reference estimates are transformed into the log2 ratio space and median normalised using the median log2 ratio excluding sex chromosomes for sex matched, mismatched and mixed estimates. These estimates are again saved back into the CNest binary file format to allow efficient extraction during the next steps of CNV detection and CNV GWAS testing.

For CNV calling CNest implements a single custom designed 3 state Hidden Markov Model (HMM) to call losses and gains, the basic implementation of which can be found (https://github.com/tf2/ViteRbi). In our hands this HMM model has been highly reliable across a number of different detection applications, and it is extremely efficient in terms of speed and memory usage. We apply a few important steps during the HMM training (using the EM algorithm) to further improve the reliability of the model. Primarily these include training the model independently for each sample by using all the log2 ratio estimates of all samples that made up the dynamic reference for that sample. By doing this we aim to increase the accuracy of the transition probabilities by giving the EM algorithm sight of large numbers of highly similar datasets. Importantly the log2 ratio estimates that we use during this training phase are those generated using the sex mismatched dynamic reference, ensuring that there is always at least one large single copy number loss/gain event present in all training datasets. Having trained the HMM for each sample independently using this approach we apply the forward backward Viterbi algorithm to call the most likely sequence of state paths across each sample independently. The result of this process is the state calls (0,1,2) for every genomic position of interest across all samples, we then apply some merging criteria to obtain both state classification and CNV regions (CNVR) across the genome. In fact, although CNV callers will often impose some complex merging criteria to account for outlier points within each called CNVR (e.g. by allowing a certain number of outlier copy number estimates within a CNVR) we are so confident in the performance of the HMM that we simply merge consecutive state calls without any additional complex merging rules. As illustrated in **Supplementary Figure 1**, this process does not result in over fragmentation of CNV regions.

### CNV merging, frequency estimation and copy number principal component analysis

Having obtained reliable copy number estimates and CNV calls across all samples we apply some cross sample merging criteria to allow us to generate merged copy number events (CNVEs) with frequency information attached. To do this we merge losses and gains separately across all samples using an iterative 50% reciprocal overlap rule, building up sets of CNVRs across all samples where all member segments (calls) within each set must share at least 50% of its boundaries (start and end positions) with at least one other segment within the set. Once we obtain full closure of the set, when no additional segment can be added to the set, we adjust the final start and end position by 80% of the inner to outer start and end positions. Finally, we calculate loss, gain and overall CNV frequency and standard errors for each CNVE resulting in a set of CNV regions across the genome with frequency estimates that can be used in some of the subsequent analyses (PCA and Association testing).

On top of the frequency estimate for all merged CNVEs we also assign frequency measures to individual bait regions where we can calculate the frequency that each bait is included within any copy number event. This gives us two different sets of CNV type and frequency datasets that can be used to perform principal component analysis (PCA) across the sample space. PCA is often used in SNP GWAS to control for population structure and other technical (or sample level) variation that is less well understood but that is important to control for during genome wide association scans. Similarly, it is important to correct for larger scale differences between copy number estimates across large datasets for CNV GWAS analysis. Often during SNP based PCA the sites get filtered or subsampled to allow efficient PCA to be performed, for CNV analysis it seems important that we can run PCA analysis using both commonly variable CNV regions and rare regions separately. This is due to an observation that when including all CNV sites in PCA, the commonly variable positions tend to suck out a lot of variation and get overrepresented in the first PCs. We used iterative PCA to perform several different types of PCA using both bait level and CNV call level copy number information stratified by frequency estimates. Overall, we find that PCA based on commonly variable positions are better able to capture sample level information such as population structure, whereas PCA based on rare regions can account for cryptic sample differences which are likely due to certain noise properties of the data that we were unable to accurately model during the previous steps.

### Genetic association testing

One major point of the CNest methods and approach is that by working with copy numbers in this way we have been able to employ genetic association testing methods similar to the SNP based GWAS methods that have been applied with great success over many years. For CNV we can use several different estimate types to perform large scale genome wide association tests. Although it would be possible to develop methods for copy number genotyping (i.e. actual copy number) due to the way we have set up our large scale approach we are not able to accurately determine the real copy number of any individual genome position. Rather we have well calibrated relative copy number estimates across large numbers of samples that can be used to search for associations against any given traits.

We set our models up in several different ways but always (in this study) by using standard linear and logistic regression techniques, although this choice is potentially suboptimal (**Discussion**) this was done deliberately to ensure that any CNV association signals follow the general additive model (where the copy number estimate must display a linear relationship against the given trait). All models were applied to unrelated samples from the PCA-defined European cluster (SNP PCs 1 and 2). For quantitative traits we use generalised linear models with covariates and use both the bait level copy number estimate (log2 ratio) and the copy number estimate (mean log2 ratio) for all merged CNVEs across all samples as the test variables. The standard set of covariates we include are sex, age, sequencing batch, the first 10 PC from SNP based PCA and the first 10 PCs from CNV PCA for both rare and common sites separately. Additionally, to ensure that outliers in the phenotype distribution do not impact our association tests, for all quantitative traits we apply an inverse rank normalisation.

For SNP based association tests in exactly the same sample sets we used bgenie (172) for quantitative and regenie for binary case control trait tests (173). Imputed SNP genotypes from the 500K UK Biobank release (172) were remapped to genome build hg38 using the UCSC liftover tool (174) prior to sample selection based on the 162,633 samples that were used in the CNV association tests. In both cases, for quantitative and qualitative tests, we followed the standard SNP filtering recommendations, including only bi-alleilc SNPs with a minor allele frequency greater than 1%. Association tests were run across the main traits of interest and the genome wide significance cut-off of 5e-08 was used to define associations between SNPs and traits.

### Definition of the association significance threshold

To justify the use of the widely accepted genome wide significance threshold of 5e-08 for significance testing in this work we looked at how our results would change when using three different p value correction approaches. We assessed the use of a stringent Bonferroni correction, the Benjamini Hochberg (BH) false discovery rate (FDR) based approach and permutation tests.

Overall, applying the stringent Bonferroni correction only slightly lowered the significance threshold to a value of 3.35e-08 and had very little effect on the number of significant exon level associations for most tests (**Supplementary Figure 7A**) with a median decrease of 0 (mean decrease of 1.19 and maximum decrease of 21) across all traits. However, applying this stringent correction did result in the exclusion of 0.5% (45/862) fine mapped regions across 18 of the 78 traits tested, after correction only 0.5% (4/78) of those traits had zero remaining significant associations at the specific loci since each of these 4 traits only had a single low level exon signal. Next, we applied a 0.01 FDR based BH correction to each association result independently, again we observed highly consistent results for most traits with a median increase of 1 significant exon signal across all traits (**Supplementary Figure 7B**). For the majority of traits, the BH correction resulted in less stringent thresholds than both the genome wide and Bonferroni approaches (**Supplementary Figure 7C**), and in most cases resulted in either the same (28 traits) or increased (40 traits) numbers of exon level signals that would be defined as significant. For some traits (8/78) the number of additional significant associations for the BH correction was substantially higher (greater than 100 additional exonic signals) suggesting that there could be some value in applying an FDR based correction. Finally, we performed 100 rounds of permutations on 4 main traits (hair colour, height, MI and Asthma) where we randomly ordered the phenotype measurements or case labels and ran our standard linear or logistic regressions models across 100 different random sets for all 4 traits. In all cases after 100 rounds of permutations there were zero signals that passed a genome wide threshold of 5e-08 (**Supplementary Figure 8**) indicating that this value is suitable for use in the definition of copy number associations.

Although it may be possible to use a less stringent threshold (such as BH or permutation based) and to obtain a greater number of copy number based trait association, we preferred to remain highly stringent. Copy number associations often display a similar association pattern to SNP GWAS tests genome wide where close by exon signals are highly correlated and associate to the same trait (LD peaks) and the use of the 5e-08 significance value had the additional benefit of allowing us to apply the same definition of significance for both SNP and CNV based association tests.

### Identifying associated genetic loci and fine mapping

We have developed a set of tools that build on top of the CNest framework to allow large-scale genome wide association testing for CNVs - CNassoc. These tools perform several important steps described above - namely CNV merge, PCA and GWAS testing using regression models. Since we have placed CNV GWAS analysis into a similar framework to that often used for SNP GWAS analysis we can make use of standard approaches for genetic loci detection and quality control. Firstly, we use the accepted genome wide significance threshold of 5e-08 to define associations between copy number and traits, although in this case we could theoretically lower this cut-off by using, for example, a FDR correction we preferred to remain highly stringent for the results we describe in this work. It also now becomes possible to use standard diagnostic approaches to association results, such as QQ plots and permutation. We apply these standard approaches to the CNV association results described here and see that in general the distributions of p values from our association tests are well controlled. For some tests we do see a degree of inflation and calculate the inflation factor - lambda - for all tests (**Supplementary Table 1**), overall for the majority of tests we get inflation factors below 1.13 which is generally considered to be acceptable in GWAS tests (175) and for cases where the inflation factor is above 1.13 we suggest that a level of caution is used when interpreting these results.

We did not perform fine mapping of SNP based association signals as it is not a focus of this work to perform SNP based GWAS, however we did fine map CNV signals to define the fine mapped regions of CNV association that we report (**GWAS catalogue**) and that we could use for investigation into CNV compared to SNP level signals during the next steps. Because our genomic test loci are, by definition, in coding regions of the genome we choose to use a relatively simple approach for fine mapping CNV associations in a gene/exon centric way. First, we merged all directly adjacent significant signals that had no intervening signal below the significance threshold, next we merged all significant signals that were found in the same gene(s). This resulted in a fine mapped list of significant copy number regions that can contain a single exon, multiple exons within a gene or multiple exons across multiple genes, to be clear these regions do not always contain the full coding region of the gene however any intervening not significant signals between two significant signals within the same gene are merged as that intervening region is assumed to be important for the underlying CNVs. For reporting purposes, the −log10 p value is reported for the lead exon signal within each fine mapped region and tests for correlation between SNP and CNV signals in the next section are always performed in relation to the lead CNV signal for each fine mapped region.

### Comparison between SNP and CNV association signals

One question that we wanted to address with this work is that of how SNP and CNV associations for the same traits interact with each other and how many of the CNV specific association results would have been detectable using standard SNP GWAS tests. To explore this we defined a set of classification rules to allow us to classify each fine mapped CNV region as either, CNV-only (not detectable using SNP GWAS), CNV-allele (detectable signal from SNP-GWAS but very hard to discover using tagging variation), SNP-CNV-near (detectable by SNP GWAS and very likely to be fine mapped to the same gene), SNP-CNV-far (detectable by SNP GWAS however likely to be mapped to a different gene).

First we calculated r^2 between the lead CNV signal and all SNP genotypes within 1 MB. If no significant SNP signal within 1MB was detected the fine-mapped region was classified as CNV-only i.e. not detectable by standard SNP GWAS. Regions are classified as CNV-allele if there were significant SNP associations within 1MB but if none of those SNPs had an r^2 greater than 0.6 i.e. association is detectable by both SNP and CNV GWAS but are not tagging. Next, if any significant SNP within 1MB did have an r^2 greater than 0.6 and if any of those SNPs were closest to the gene(s) inside the fine mapped CNV region than the region was classified as SNP-CNV-near i.e. the association signal was taggable by a SNP that associated with the same trait and was highly likely to be assigned to the same gene. Finally, if there were significant SNPs within 1MB that have an r^2 greater than 0.6 but if all those SNPs were closer to a gene that was not within the fine-mapped CNV region then we classify these as SNP-CNV-far. It is worth noting that we set our SNP to CNV r^2 cut-off quite low at a value of 0.6, this is because genome wide we observe relatively low r^2 between SNPs and CNVs and although there are numerous cases of very well tagged CNVs (r^2 > 0.9) for these results we decided to be strict with the definition of novel CNV associations meaning that when we classify an associated fine-mapped CNV region as CNV-allele we can be very confident that it is not well tagged by any associating SNP within 1 MB and this results in an overall decrease in the number of CNV associations that we classify as CNV-allele.

### Competitive SNP-CNV association models

To look in greater detail at the relationship between SNP and CNV association interactions we performed some joint modelling of two different copy number estimates, log2 ratios and approximated copy number state distributions, by including either the SNP genotypes from the most significant SNP association or the best tagging SNP (highest r^2 values to the lead CNV association) within 1MB around the CNV association location as covariates within a standard linear model. First we selected 91 single exon associations across 2 traits (hair colour and standing height) and extracted the log2 ratio values across all UK Biobank samples included in the association testing and fitted a 3 component mixture model to define the approximate copy number states boundaries, the decision to use a 3 component was two fold, firstly it was to avoid complications in the definition of the number of actual copy number states observed across the 91 sites that would be highly likely to cause problems during mixture model fits and secondly it was to place the copy number state models into a similar categorical distribution to the SNP genotype models. Each sample was assigned a copy number state based on its most likely component from the mixture model and additionally we did not allow any sample to cross the mean of any adjacent component resulting in a 3 state copy number model relating to low, medium and high copy numbers. Next, we fitted 8 different types of models across all 91 sites where we tested all variant classes independently and additionally included both SNP types (most significant association and best tagging SNP within 1MB around the CNV association) in pairwise competitive linear models for the copy number estimate and copy number state distributions. We extracted both p values and beta effect sizes from each variant type from all the eight different models and carried out a comparison of signal strength across the different models, allowing us to look in more detail at the relationship and interactions between SNP and CNV associations using close to one hundred CNV association discoveries.

## Code availability

The main CNest code base and docker setup can be found here: https://github.com/tf2/CNest This repository contains all the source code and a docker setup as well as NextFlow and WDL workflows. We are in the process of developing a detailed tutorial for getting CNest up and running across a diverse set of computational infrastructure including cloud based systems. This tutorial and the all additional workflow implementations will be linked to from the main CNest repository as soon as available.

## Data availability

CNV GWAS summary statistics and fine mapped association regions are included in the supplementary material of this paper. CNV calls and copy number estimates will be made available via the UK BioBank data return and linked to UK Biobank application number: 49978. Additionally, CNV-GWAS results will be uploaded into an appropriate portal within the GWAS catalogue.

## Acknowledgements

We acknowledge all the UK BioBank participants and the UK Biobank management team without whom this study would not have been possible. T. Fitzgerald and E. Birney were funded by the EMBL European Bioinformatics Institute (EMBL-EBI).

## Author contributions

T.F and E.B conceived and designed the study, performed the analysis, and wrote the manuscript.

## Competing interests

The authors declare no competing interests.

## Notes

### Competing Interest Statement

The authors have declared no competing interest.

