## Supplementary figures for "CNest: A Novel Copy Number Association Discovery Method Uncovers 862 New Associations from 200,629 Whole Exome Sequence Datasets in the UK Biobank"

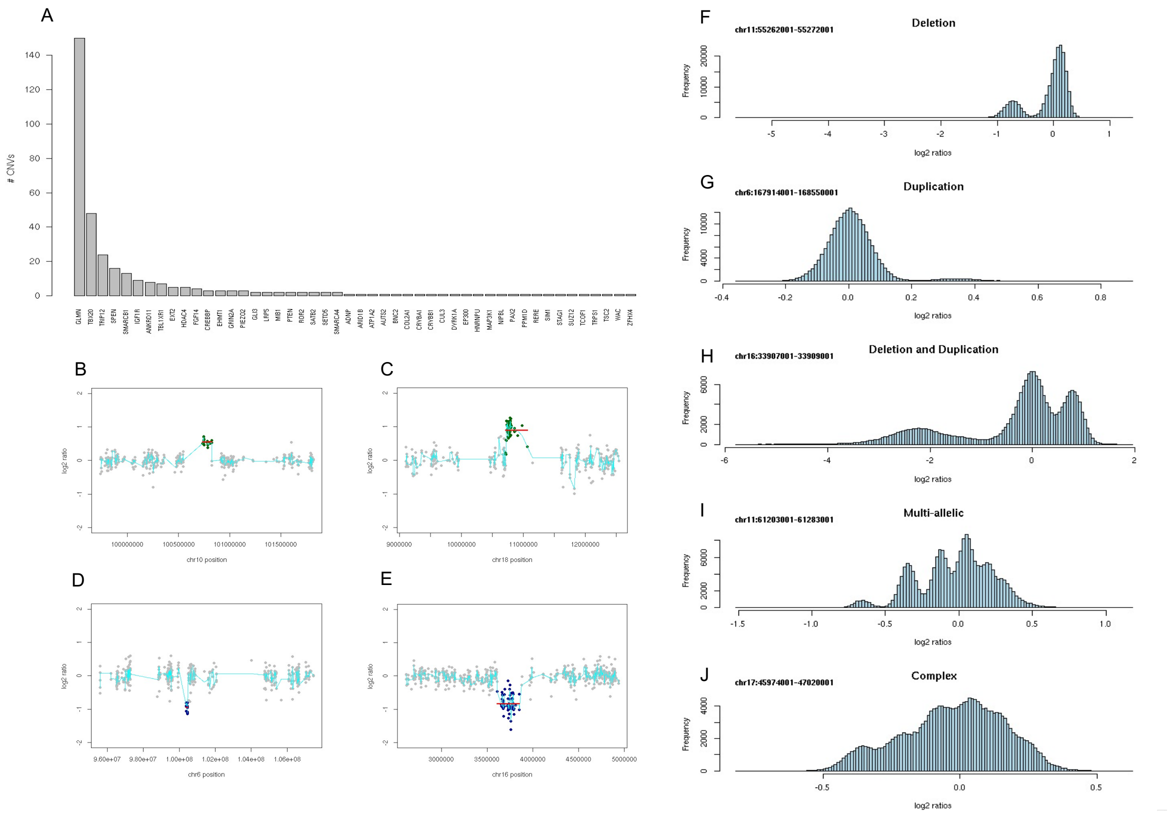


*Supplementary Figure 1: Individual CNV calls and copy number variable locations in the UK Biobank. A: Barplot showing the number of CNV calls overlapping any of 218 monoallelic loss of function genes from the DDG2P (dd gene to phenotype). B: Truncating duplication at the PAX2 gene. C: Truncting duplication at the PIEZO2 gene. D: Deletion at the SIM1 gene. E: Deletion at the CREBBP gene. F: Deletion locus at 11q12.1. G: Duplication locus at 6q27. H: Deletion / duplication locus at 16p11.2. I: Multi-allelic locus at 11q12.2. J: Complex locus at 17q21.31.*

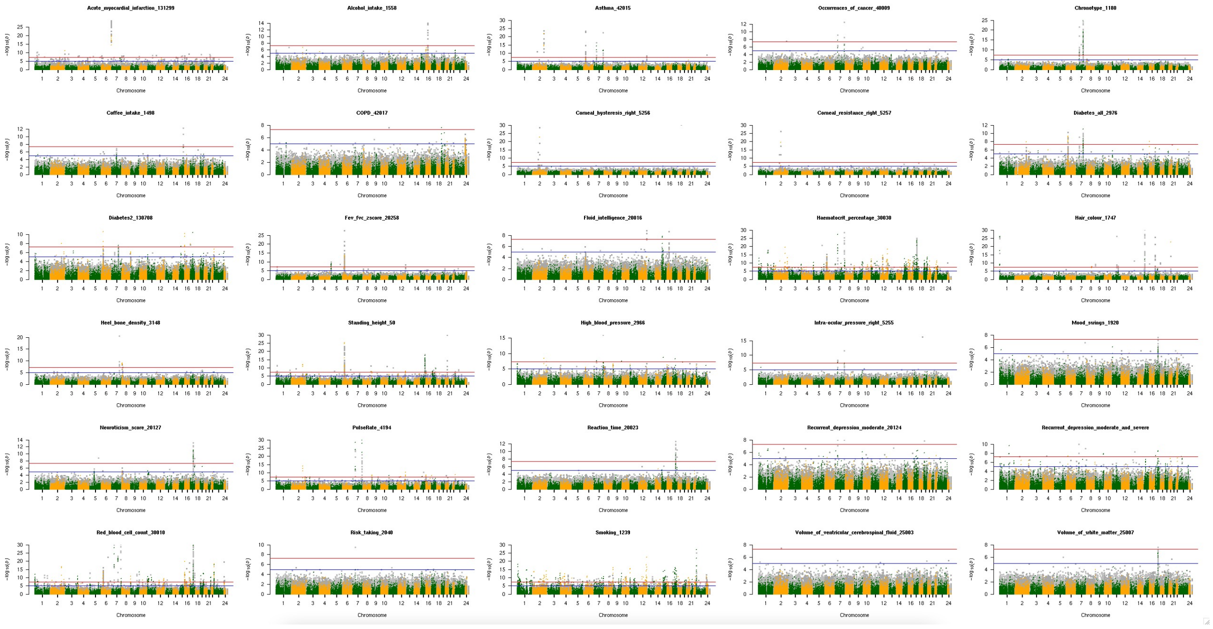

Supplementary Figure 2*: Individual manhattan plots for 30 of the 34 main traits for CNV association in the UK Biobank 200K whole exomes. All plots are pinned to a maximum -log10 p-value of less than 30, meaning that all stronger association signals are not shown but this significantly aids the visualisation across all traits. We exclude 4 traits, showing only right eyes for eye related traits and only red blood cell counts for red blood cell related traits. All of the 30 panels have a title showing the trait and all include both p-values from exon level (copy number estimates) trait association testing (grey) and CNV call association testing (green and orange).*

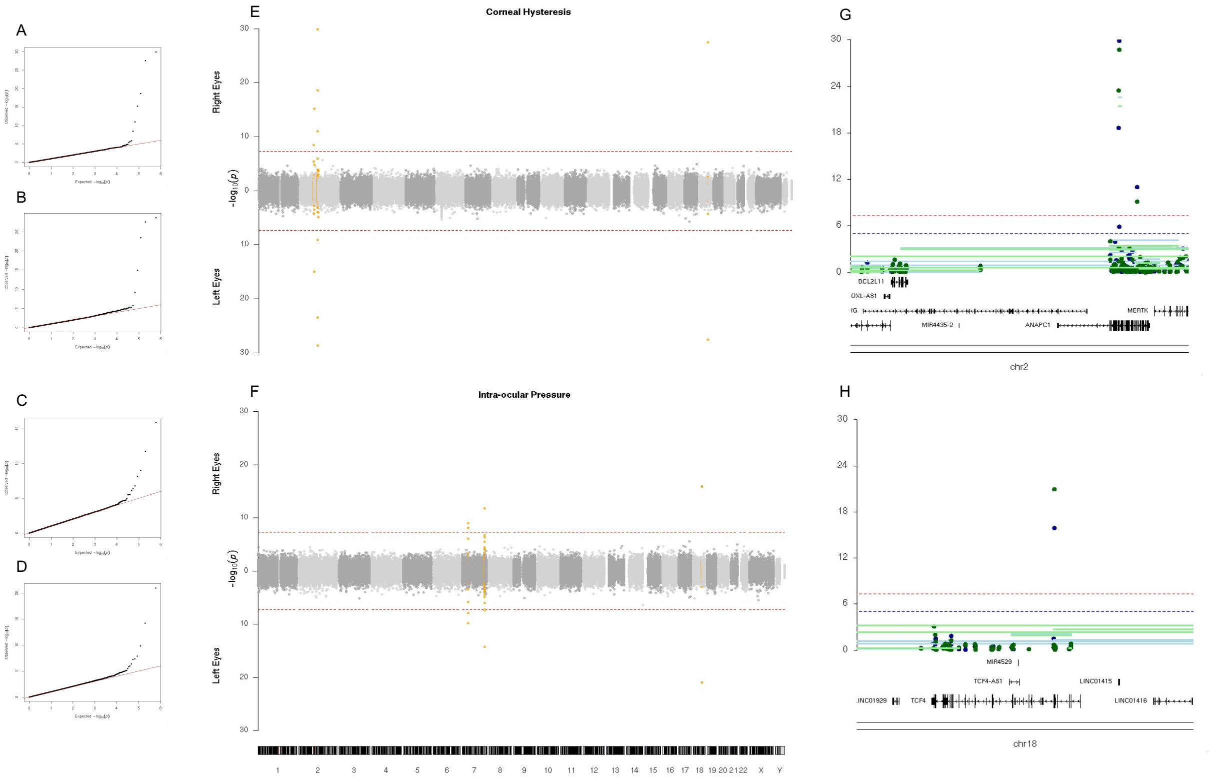


*Supplementary* Figure 3: *CNV association results for the eye related traits, Corneal hysteresis and Intra-ocular pressure for left and right eyes separately. A: QQ plot for Corneal hysteresis in right eyes. B: QQ plot for Corneal hysteresis in left eyes. C: QQ plot for Intra-ocular pressure in right eyes. D: QQ plot for Intra-ocular pressure in left eyes. E: Bidirectional manhattan plot for Corneal hysteresis in right (top) and left (bottom) eyes. F: Bidirectional manhattan plot for Intra-ocular pressure in right (top) and left (bottom) eyes. G: Locus zoom plot of right eye Corneal hysteresis at the ANAPC1 gene. H: Locus zoom plot of right eye Intra-ocular pressure at the TCF4 gene.*

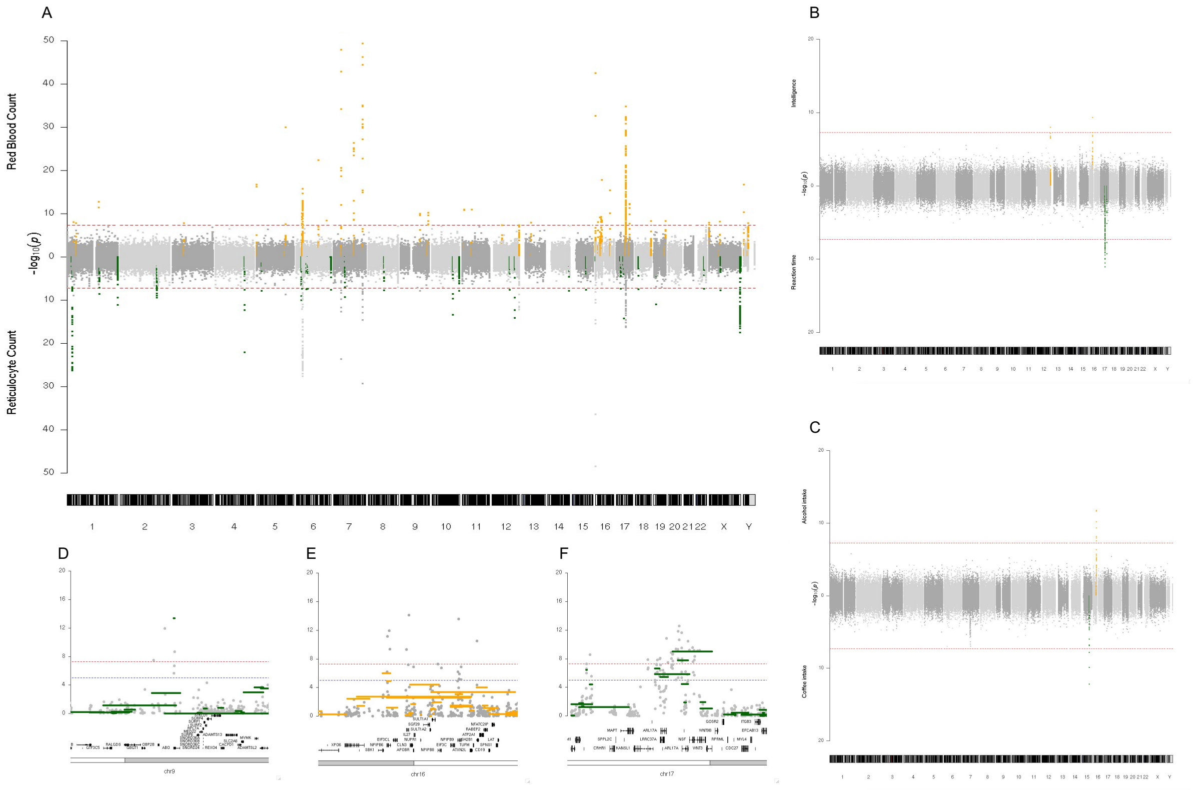


Supplementary *Figure 4: CNV association results for the red blood cell, neurological and behavioural related traits. A: Bidirectional manhattan plots for red blood cell count (top) and reticulocyte count (bottom), fine mapped regions are highlight in orange (red blood cell count) and green (reticulocyte count) with fine mapped regions for reticulocyte count that were also discovered for red blood cell counts not being highlighted. B: Bidirectional manhattan plots for fluid intelligence (top) and reaction time (bottom), fine mapped regions are highlighted in orange and green respectively. C: Bidirectional manhattan plots for alcohol (top) and coffee (bottom) intake, fine mapped regions are highlighted in orange and green respectively. D: Locus zoom plot for red blood cell count at the ABO gene. E: Locus zoom plot for alcohol intake around the NPIPB6 gene. F: Locus zoom plot for reaction time around the ARL17B gene.*

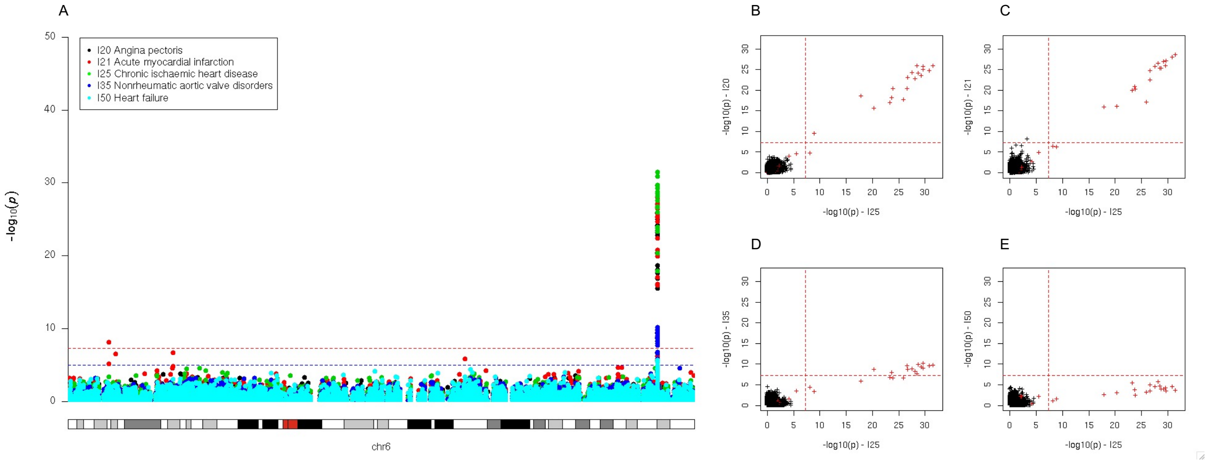


*Supplementary* Figure 5: *Comparison of association signal strength for heart related ICD10 codes at the LPA gene. A: Overlaid manhattan plot from chromosome 6 including 5 heart related ICD10 code based case/control tests. B: Minus log10 p values for ICD10 code I25 against I20. C: Minus log10 p values for ICD10 code I25 against I21. D: Minus log10 p values for ICD10 code I25 against I35. E: Minus log10 p values for ICD10 code I25 against I50. The points highlighted in red in panels B - E are all exons within the LPA gene.*

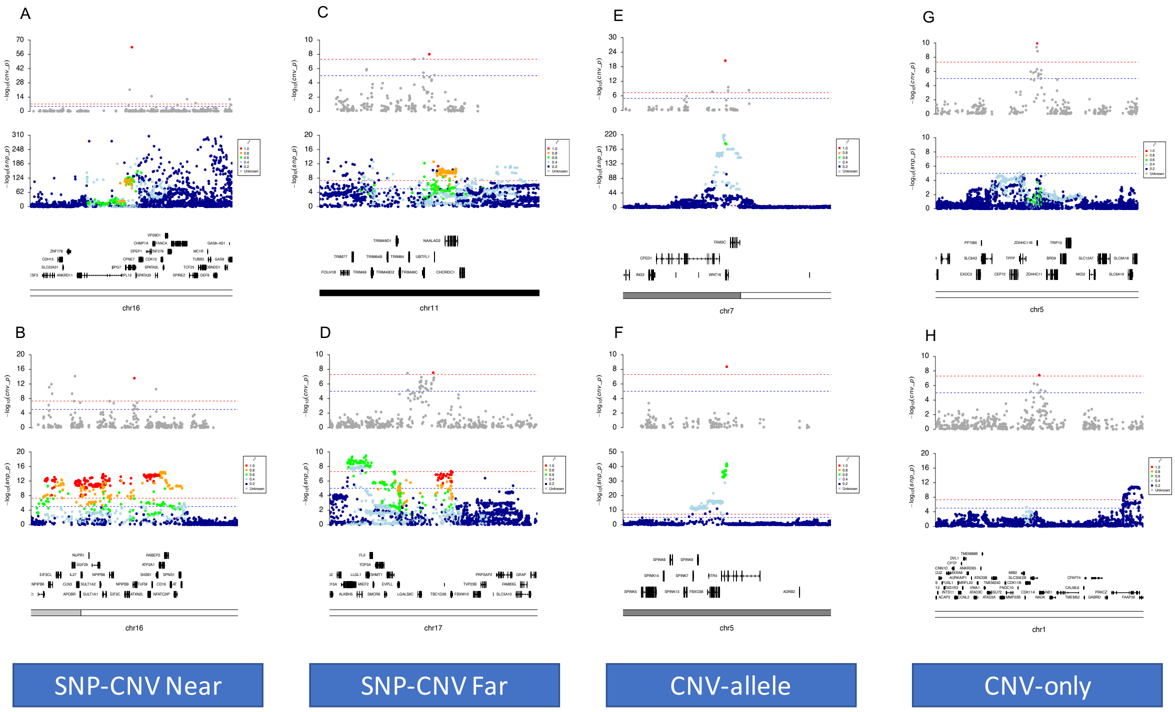


Supplementary *Figure 6: Locus zoom plots showing some CNV association categories. A: SNP-CNV near discovery for hair colour involving exons 7-10 of the SPG7 gene. B: SNP-CNV near discovery for alcohol consumption at a 0.8MB region containing 4 fine mapped CNV regions involving the NPIPB6, NPIPB7, NPIPB9 and SH2B1 genes. C: SNP-CNV far discovery for hair colour at the TRIM49C with tagging SNPs downstream at UBTFL1 or NAALAD2 genes. D: SNP-CNV far discovery for standing height at a 12.6KB region including the EVPLL and LGALS9C genes. E: CNV-allele discovery for heel bone density at the WNT16 gene. F: CNV-allele discovery for the FEV/FEC ratio involving exon 1 of the HTR4 gene. G: CNV-only association for the FEV/FEC ratio on chromosome 5 at the ZDHHC11B gene. H: CNV-only discovery for standing height including several genes that are pulled up towards suggestive genome wide significance with a single exon signal that passes genome wide significance within the CDK11A gene.*

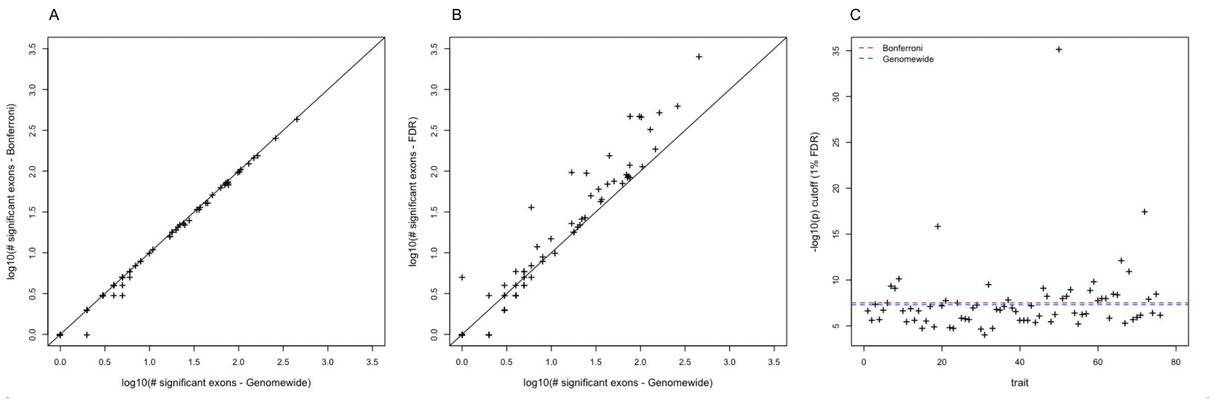


Supplementary *Figure 7*: *Comparison of significance level approaches. A: log10 of the number of significant exon level signals per trait using the genome wide 5e-08 cut-off vs. a strict Bonferroni cut-off. B: log10 of the number of significant exon level signals per trait using the genome wide 5e-08 cut-off vs. a 1% FDR cut-off. C: The -log10 significance level at a 1% FDR cut-off for all traits tested. The blue dashed line indicates the genome wide 5e-08 cut-off and the red dashed line indicates the Bonferroni cut-off.*

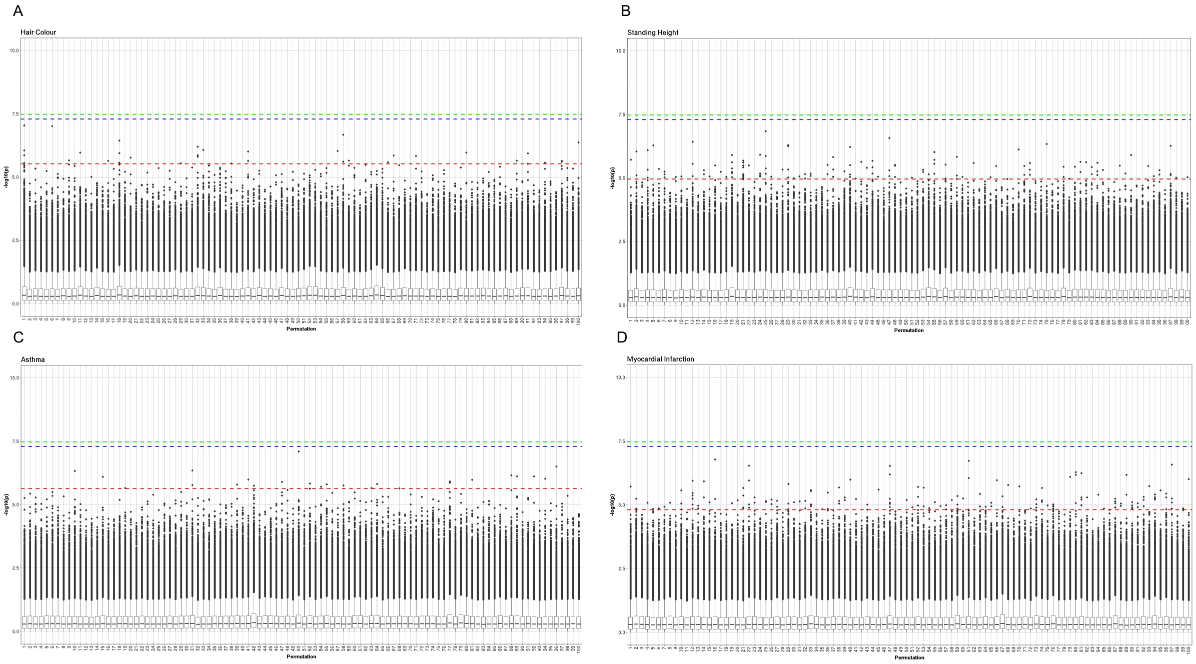


Supplementary *Figure 8: Permutation tests for the 4 main traits. Tests were performed using 100 different random ordering for each trait or case label followed by association testing genome wide. A: Genome wide tests across 100 differently permuted datasets for hair colour. B: Genome wide tests across 100 differently permuted datasets for standing height. C: Genome wide tests across 100 differently permuted case labels for asthma. D: Genome wide tests across 100 differently permuted case labels for myocardial infarction. The red dashed line indicates the 1% FDR cut-off, the blue dashed line indicates the genome wide 5e-08 cut-off and the green dashed line indicates the Bonferroni cut-off.*
